# A genome-wide CRISPR-Cas9 screen identifies CENPJ as a host regulator of altered microtubule organization during *Plasmodium* liver infection

**DOI:** 10.1101/2020.08.31.275867

**Authors:** Kamalakannan Vijayan, Nadia Arang, Ling Wei, Robert Morrison, Rechel Geiger, K. Rachael Parks, Adam J Lewis, Fred D Mast, Alyse N Douglass, Heather S Kain, John D Aitchison, Jarrod S Johnson, Alan Aderem, Alexis Kaushansky

## Abstract

Prior to initiating symptomatic malaria, a single *Plasmodium* sporozoite infects a hepatocyte and develops into thousands of merozoites, in part by scavenging host resources. We show that host microtubules dynamically reorganize around the developing liver stage (LS) parasite. Using a genome-wide CRISPR-Cas9 screen, we identified host regulators of cytoskeleton organization, vesicle trafficking, ER/Golgi stress and lipid biogenesis that regulate *Plasmodium* LS development. These novel regulators of infection, including Centromere Protein J (CENPJ), led us to interrogate how microtubule organizing centers (MTOCs) are regulated during infection. Foci of γ-tubulin localized to the parasite periphery; depletion of CENPJ exacerbated this re-localization and increased infection. Further, we show that the Golgi acts as a non-centrosomal MTOC by organizing γ-tubulin and stimulating microtubule nucleation at the parasite periphery. Collectively, we show that the *Plasmodium* LS recruits the host Golgi to form MT mediated conduits along which host organelles are recruited to the PVM, to support liver stage development. Our findings suggest many host-targeted pharmacological inhibitors may inhibit LS infection.

## Introduction

Malaria is transmitted to humans by the injection of *Plasmodium* sporozoites into the skin during the blood meal of an infectious female *Anopheles* mosquito. Sporozoites exit the skin by traversing blood vessels to enter the circulation and are then carried to the liver. Sporozoites leave the circulation by traversing the sinusoidal cell layer and infecting hepatocytes and forming liver stages (LS) (Mota et al., 2001; Shortt and Garnham, 1948; Vanderberg, 1981). LS parasites reside in a membrane-bound compartment in the hepatocyte termed the parasitophorous vacuole (PV), where they differentiate into exoerythrocytic merozoites. The PV membrane subsequently breaks down, and merozoites reenter the blood and infect erythrocytes (Sturm et al., 2006).

Hepatocyte infection is obligate for parasite life cycle progression, and thus is an important target for antimalarial intervention. Elimination of the parasite during this stage would block both disease symptoms and transmission. During LS development, host hepatocytes undergo remarkable morphological changes. The cytoskeleton is a key regulator of deformability, and *Plasmodium* has been demonstrated to alter host cell actin cytoskeleton during LS development (Gomes-Santos et al., 2012), egress (Burda et al., 2017), and blood stage development (Hale et al., 2017; Warncke and Beck, 2019). We and others previously have demonstrated that the *Plasmodium* liver stage PV membrane interacts with late endosomes (Petersen et al., 2017), lysosomes (Lopes da Silva et al., 2012; Niklaus et al., 2019; Prado et al., 2015; Risco-Castillo et al., 2015; Vijayan et al., 2019), retrograde vesicles (Raphemot et al., 2019) and autophagic vesicles (Prado et al., 2015; Real et al., 2018; Wacker et al., 2017). As many intracellular pathogens target the host microtubule network to subvert host vesicle trafficking events for their own benefit (Alix et al., 2011; Asrat et al., 2014), we hypothesized that *Plasmodium* LS parasites actively alters the host cytoskeleton to traffic the host vesicles to PVM.

Multiple focused forward genetic screens have informed our understanding of host regulatory factors for LS malaria (Prudencio et al., 2008; Raphemot et al., 2019; Rodrigues et al., 2008). These screens have provided valuable insights into parasite-host interactions, but the scope of these investigations have been limited, suggesting that a complete complement of factors required for *Plasmodium* entry and development remains to be discovered. *Plasmodium* LS infection actively remodels the host hepatocyte by rewiring a portion of host cell signaling and disrupting canonical signaling cascades (Glennon et al., 2019). We therefore sought to use an unbiased genome-wide approach to identify the parasite driven host factors that contribute to host cytoskeleton remodeling.

Recently, genome-wide CRISPR screening has emerged as a powerful strategy to identify novel gene functions (Sanjana et al., 2014; Shalem et al., 2014). We report the first genome wide CRISPR-Cas9 screen that aims to identify host factors that regulate *Plasmodium* infection. The screen identified several host factors critical for *Plasmodium* development, comprising previously explored and novel host regulators, including those that lead to rearrangement of the cytoskeleton and vesicular trafficking. Our work provides the foundation for a more comprehensive understanding of the host processes that are required for optimal *Plasmodium* development and could aid in the efforts to develop host-targeted antimalarial drugs.

## Results

### *P. yoelii*-infected cells exhibit alteration in microtubule organization

To visualize microtubule (MT) organization, we transfected HepG2-CD81 cells with CellLight™ rfp-α-tubulin BacMam 2.0 and infected the cells with *P. yoelii* sporozoites for 6 and 24 h (Fig. 1A). Strikingly, we observed that the host MT network redistributes to the LS parasite, appearing to wrap around the PVM at 24 h.p.i but not at 6 h.p.i (Fig. 1A and B). In contrast, MTs in uninfected cells form a canonical network around the nucleus, radiating toward the cell periphery (Fig. 1A). Acetylated MT are the stabilized form of MT that support kinesin-mediated trafficking of vesicles (Reed et al., 2006). We next asked if the MTs associated with the parasite are actively engaged in cell transport by assessing their acetylation status. We infected HepG2-CD81 cells with *P. yoelii* sporozoites, and then visualized acetylated alpha-tubulin and the parasite PVM-resident protein UIS4 by immunostaining (Fig. 1C). MTs that decorate the parasite periphery were highly acetylated. In contrast, in uninfected cells, acetylated MTs were distributed throughout the cytosol (Fig. 1C). We next disassembled the MT by nocodazole treatment (Zhu and Kaverina, 2011). Briefly, we infected rfp-α-tubulin transfected HepG2-CD81 cells with *P. yoelii* sporozoites, and allowed the infection to procced for 48 h. After 46 h, nocodazole was added. After an additional 2h (48h post-infection), cells were washed, and incubated with nocodazole-free media for 45 sec to allow the nucleation of MT(Zhu and Kaverina, 2011). In uninfected cells, microtubules nucleated close to host nucleus (Fig. 1D). Strikingly, in the infected cells, MTs nucleated adjacent to parasite periphery. Together, these results suggest that during LS development, *P. yoelii* remodels the host MT network to nucleates from the parasite periphery.

**Figure 1.**
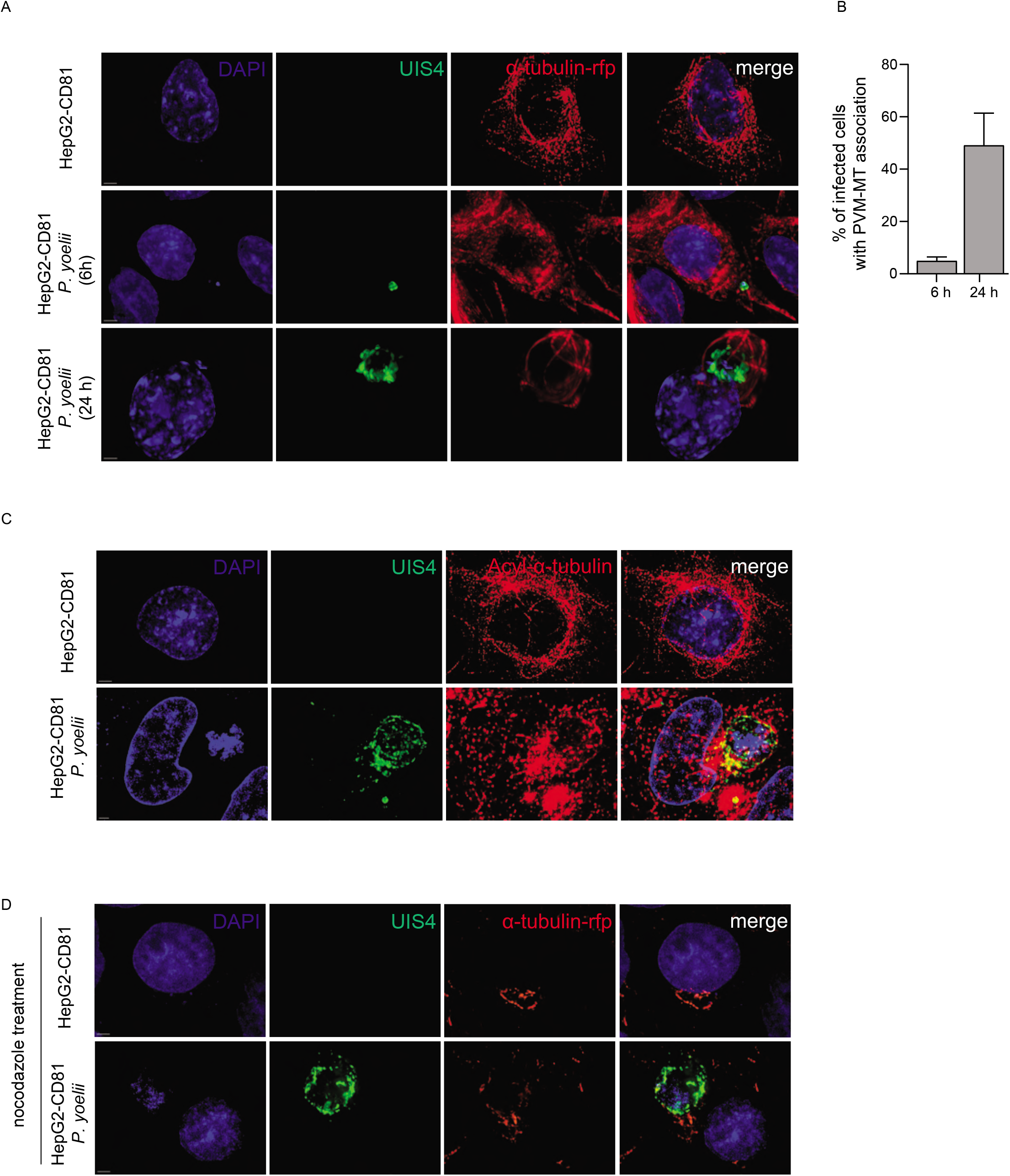
*P. yoelii*-infected cells exhibit alterations in microtubule organization. **(A)** HepG2-CD81 cells transfected with CellLight™ rfp-α-tubulin BacMam 2.0 were infected with *P. yoelii* sporozoites for 6 and 24 h. Cells were fixed and stained with antibodies to UIS4. Images are maximum intensity projections from 20-25 z-slices. **(B)** The bar graph represents parasitophorous vacuolar membrane (PVM) and MT association. Images and the bar graph are a representative of 3 independent experiments with n ≥ 100 cells / condition. **(C)** HepG2-CD81 cells were infected with *P. yoelii* sporozoites, fixed and stained with antibodies to *P. yoelii* UIS4 and acetylated α-tubulin. Images are maximum intensity projections from 20-25 z-slices. **(D)** HepG2-CD81 cells were infected with *P. yoelii* sporozoites for 48 h. Nocodazole was added to the cells after 46h for 2 hours. Cells were washed and incubated for 45 sec to allow the nucleation of MT. Cells were fixed and stained with antibodies to UIS4-647 conjugate. Images and bar graph are a representative of 3 independent experiments with n ≥ 100 cells / condition. Bar = 2 μm.

### CRISPR-Cas9 screen to identify host regulators of infection

Whole genome CRISPR-Cas9 screens facilitate an unbiased approach to identify regulators of a key process. To identify host genes critical for MT remodeling during *Plasmodium* LS development, we prioritized breadth in our approach, as canonical regulators of infection are sometimes rewired during infection (Glennon et al., 2019). We used a pooled library of GeCKOv2 sgRNAs to generate a whole-genome knockout library in HepG2-CD81 cells (Sanjana et al., 2014; Shalem et al., 2014). HepG2-CD81 cells were transduced with lentivirus containing the pooled GeCKOv2 sgRNA library of 123,642 sgRNAs targeting 19,031 protein-coding genes (~6 sgRNAs/gene), 1,864 microRNAs (4 sgRNA/microRNA) and 1,000 negative controls (2 sgRNA/control) and selected in puromycin for 5-7 days. To evaluate sgRNA diversity in the HepG2-CD81-GeCKOv2 library, we PCR-amplified the integrated sgRNA cassettes from genomic DNA extracted from transduced cells and subjected the amplified library to Illumina sequencing. At the gene level, 16,629 out of 19,031 (87.38%) genes targeted by 3 or more sgRNAs guides were significantly enriched. We observed an absence of sgRNAs targeting 2402 genes out of 19031 (12.62%); this may be due to gene essentiality or the failure of certain sgRNA to incorporate successfully into the genome. We infected forty million puromycin-resistant cells with green fluorescent protein (GFP) expressing-*Plasmodium yoelii* at a multiplicity of infection (MOI) of 0.3. After 24 h of infection, cells were sorted into infected and bystander cell populations by GFP signal intensity with fluorescence-activated cell sorting (FACS) (Fig. 2A). Separately, a parallel culture of uninfected cells was also maintained to normalize the sgRNA frequency distributions. We obtained four independent biological replicates with library generation and sequencing occurring in parallel. Genes with significantly enriched sgRNAs were identified for both the bystander and infected populations compared to uninfected cells.

**Figure 2.**
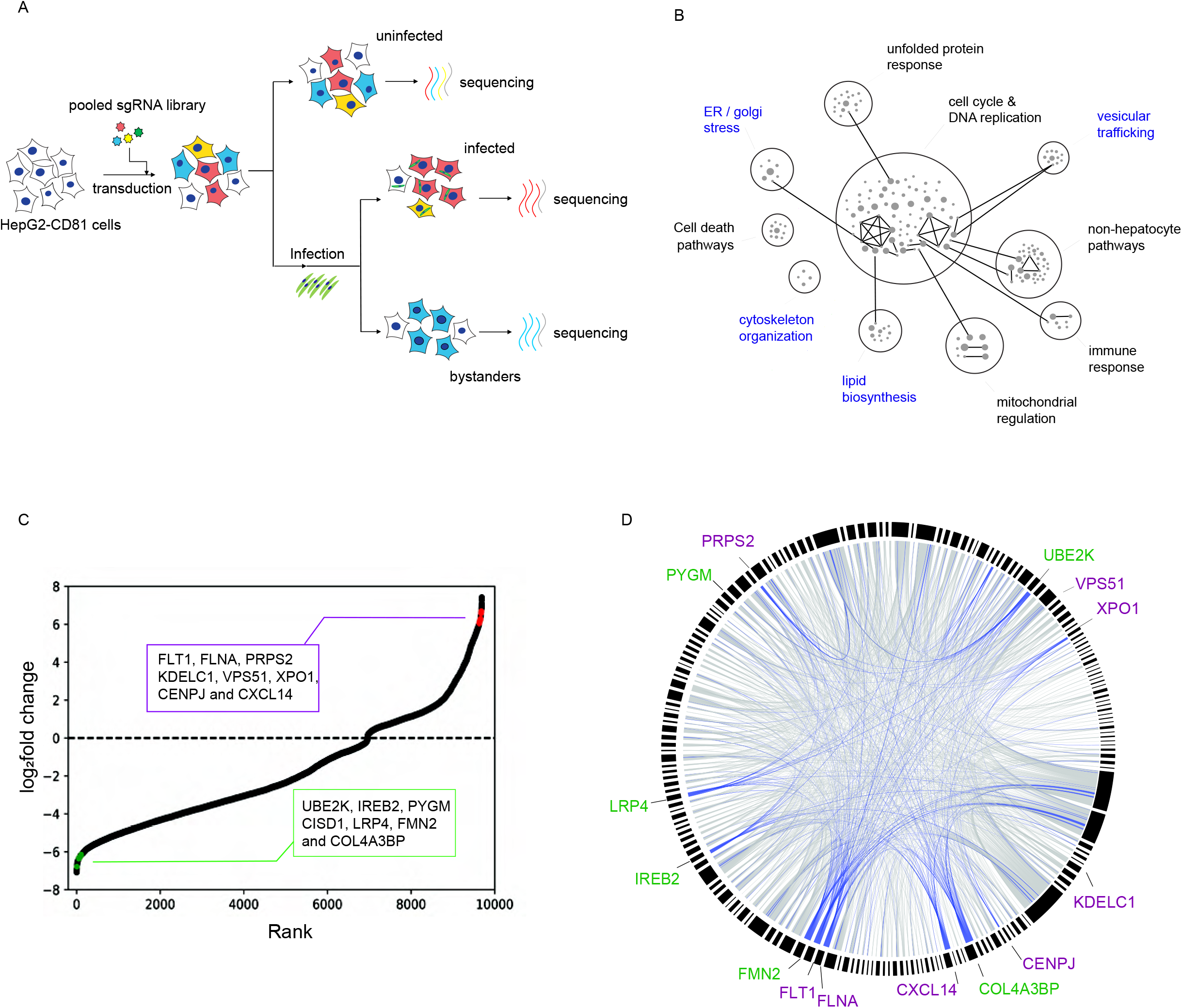
Whole genome hepatocyte CRISPR-Cas9 screen reveals putative *Plasmodium* liver stage regulatory factors. **(A)** Workflow of the hepatocyte CRISPR-Cas9 screen. A pooled CRISPR-Cas9 lentiviral sgRNA library is used to transduce HepG2-CD81 cells. Cells are infected with GFP-expressing *P. yoelli* sporozoites, and infected cells are isolated via FACS. Four biological samples were collected, and Illumina sequenced to quantify sgRNA counts from: uninfected cells, infected cells, and bystander cells. **(B)** sgRNAs observed at different levels in infected and uninfected cells are enriched in multiple GO biological processes. Nodes represent different biological processes, and the size of each node is scaled to the number of sgRNAs in the underlying gene set. Connection between nodes indicates that they share at least one gene. The nodes are grouped and further annotated. The nodes highlighted in blue are highly represented and characterized further in the study. GO terms are clustered based on higher order hierarchy using ClueGo cytoscape plugin. **(C)** Ranked log_2_ FC of genes with different levels of sgRNAs in infected and uninfected cells with a p-value < 0.05. Genes selected for further study are colored magenta (log_2_ FC > 0; negative regulators of infection) or green (log_2_FC < 0; positive regulators of infection). **(D)** A chord diagram of genes with significantly enriched sgRNAs in infected vs uninfected cells (p-value < 0.05, log_2_FC < −6.0 or > 6.0). An edge connecting genes indicates that they both belong to at least one gene set according to GO terms. Genes selected for further study are colored red (log_2_ fold change>0) or green (log_2_FC <0). Blue coded edges represent the connection between the genes belonging to the GO terms highlighted in **(B)**.

Cells that harbor genetic alterations restricting *P. yoelii* development (i.e., sgRNAs that target host genes important for infection) were expected to be enriched in the uninfected bin; we termed this group ‘putative positive regulators of infection’. We categorized sgRNAs enriched in the infected cells as ‘putative negative regulators of infection’. In this initial screen, we identified 242 genes that were statistically enriched in infected or bystander groups after accounting for multiple hypotheses. There were 67 genes significantly enriched in the infected cells compared to uninfected cells and 175 genes were significantly enriched in the bystander bin relative to uninfected bin.

To further down-select the high confidence genes, we reasoned that biological pathways with multiple putative regulators were more likely to be bona fide regulators of infection. Hence, we performed gene ontology (GO) pathway analysis to identify significantly enriched biological processes (Fig. 2B) using 242 hits from our initial screen. We shortlisted the significant gene regulators present in statistically enriched biological processes for further validation. After this stringent down-selection step, we were left with eight putative negative regulators of infection and seven putative positive regulators of infection for further study (Fig, 2C and D).

### Integrating multiple forward genetic screens provides additional testable hypotheses

We report the first global screen for host factors that regulate *Plasmodium* LS infection. Yet, like previous focused screens, our screen includes false negatives as sgRNAs are lost during the generation of the library and/or not all sgRNAs result in the disruption of the functional protein. To generate a more comprehensive picture, we systematically compared our screen, which interrogated host regulators of *P. yoelii* infection, with earlier forward genetic screens (Prudencio et al., 2008; Raphemot et al., 2019; Rodrigues et al., 2008) that identified regulators of the closely related parasite, *P. berghei* (Fig. S1A, Supplementary file 1). For the purpose of analysis, we pooled results from the screens by Rodrigues et al., 2008 and Prudencio et al., 2008, as these two screens used the same methodology but had no overlapping factors. To compare our findings to previous screens, we developed a new methodology, meta-analysis by information content (MAIC), to combine data from diverse sources, in the form of ranked gene lists. Briefly, a meta ranking of the 3 screens was performed by sorting each screen separately by z score, calculating each gene’s rank percentile location after sorting, and then averaging the gene rank percentile locations across the 3 screens, with no penalty for a gene being missing in a screen. This meta ranking was then sorted by average rank percentile location and augmented with the average z score from all screens where the function of the gene was evaluated (Supplementary file 1). Positively and negatively represented genes were sorted separately, then combined afterwards for pathway analysis. At gene level, using the z score cutoff of 2 and 1.5, our screen shared only few hits each of the other screens (Fig. S1B and S1C) without any common hits across all screens. We reasoned this could be due to many factors including host cell type, parasite species and the methodology employed. When we loosened the stringency of the cutoff to a Z-score of 1, there were several genes overlapping between the three screens, although false positive rates could be higher at this cut-off.

Despite little specific gene overlap, we asked if overlapping pathways and biological processes were present in all the screens. We employed ClueGO to determine gene ontology (GO) (Bindea et al., 2009) and observed significant enrichment in biological processes from the genes represented in at least two of three screens at z-score of 1.5 and 1. Specifically, we identified 19 high confident biological processes that are significantly enriched using a z-score cutoff of 1.5. This includes biological processes that have been previously described, such as scavenger receptor activity and cholesterol biosynthesis reported (Itoe et al., 2014; Labaied et al., 2011; Petersen et al., 2017; Rodrigues et al., 2008) (Fig. S1D and S1E). Taken together, this combined resource provides a wealth of hypotheses for further investigation.

### Identifying host factors that regulate *Plasmodium* LS invasion and development from CRISPR-Cas9 screen

To evaluate the false positive rate of our screen, we individually disrupted each of the 15 putative regulators with three sgRNAs per gene using CRISPR-Cas9 gene editing of HepG2-CD81 cells. In this system, a fluorescent reporter, GFP, is expressed only upon guide integration and puromycin resistance, enabling us to exclude any cells that did not take up and integrate the sgRNA (Fig. S2A). GFP positive cells were FACS sorted, cultured with puromycin and the knockout efficiency of CENPJ and COL4A3BP was further confirmed using western blot (Fig. S2B and S2C). To identify genes that alter *Plasmodium* LS invasion, we infected each knockout line with *P. yoelii* sporozoites for 90 min and assessed hepatocyte entry by flow cytometry. Among the selected 15 hits, only low-density lipoprotein receptor-related protein 4 (LRP4) exhibited significantly reduced entry of sporozoites 90 min after infection (Fig. 3A). This is consistent with the previous finding that CSP interacts with LRP and HSPG to facilitates host cell invasion of *Plasmodium* (Shakibaei and Frevert, 1996).

**Figure 3.**
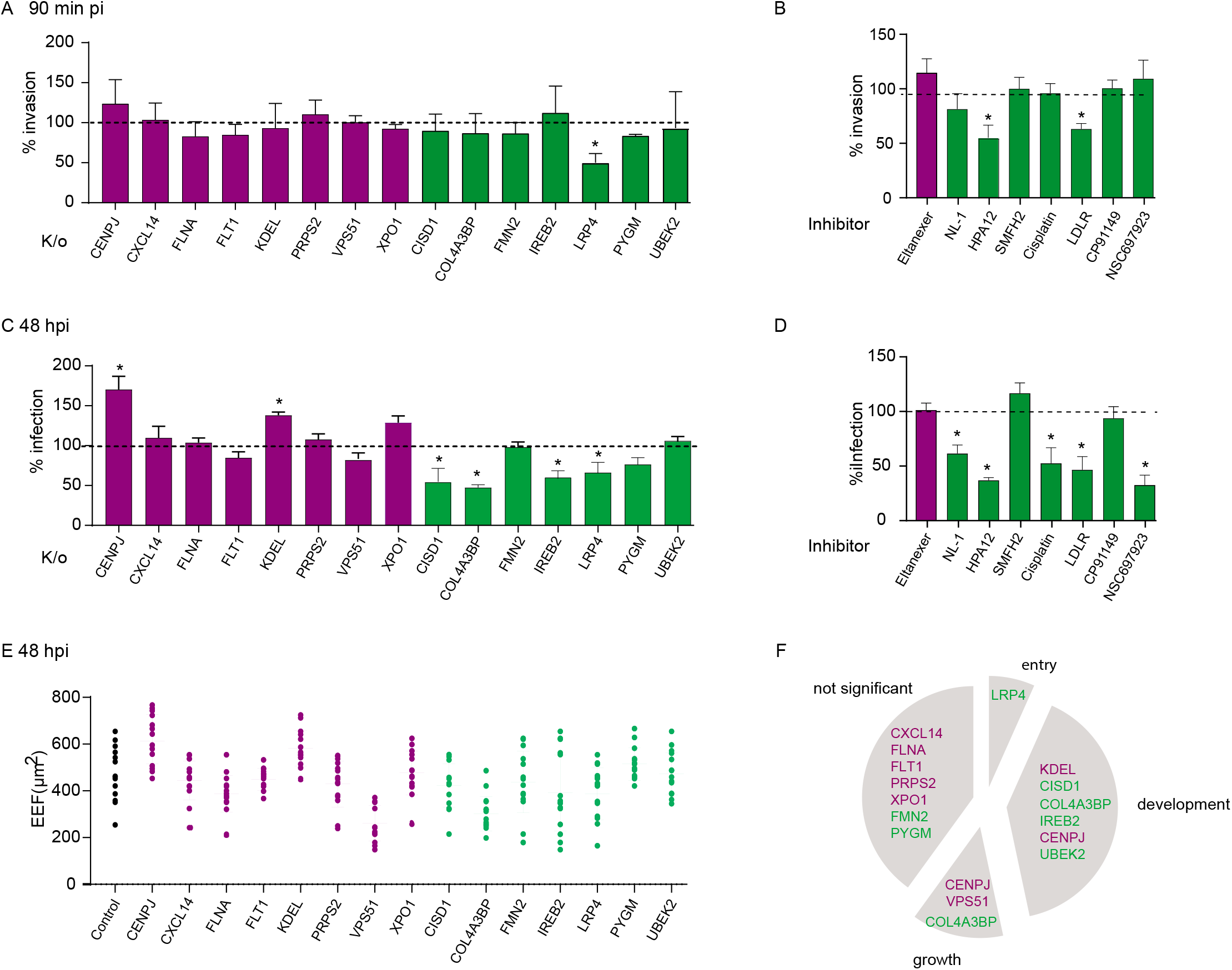
Evaluation of selected hits from CRISPR-Cas9 screen for activity in LS infection and development. **(A)** HepG2-CD81 cells were transfected with CRISPR-Cas9 containing plasmids targeting the specified gene or scrambled control and challenged with *P. yoelii* sporozoites for 90 min. The bar graph depicts invasion as the rate of *Py*CSP-positive cells for each sgRNA knockout, normalized to a scrambled control. n=3; mean ± SD. Magenta represents negative regulators of infection and green represents positive regulators of infection. **(B)** HepG2-CD81 cells were pre-treated with or without the presence of specified compounds for 2 h and infected with *P. yoelii* sporozoites for 90 min. As in (A), the bar graph depicts the invasion rate. n=3; mean ± SD. **(C)** HepG2-CD81 cells were transfected with CRISPR-Cas9 containing plasmids targeting the specified gene or scrambled control and challenged with *P. yoelii* sporozoites. After 48 h, infection was evaluated using fluorescence microscopy. The bar graph depicts the infection rate after knockout of each transcript of interest normalized to scramble cells. n=3; mean ± SD. **(D)** HepG2-CD81 cells were infected with *P. yoelii* sporozoites for 90 min, washed and treated with or without the presence of specified compounds for 48 h. As in **(C)** the bar graph depicts the infection rate. n=3; mean ± SD **(E)** Assessment the liverstage (LS) parasite size from microscopic from **(C)**. (n=3; mean ± SD). **(F)** A pie chart depicts HepG2-CD81 knockouts with significantly different entry, development or growth rates than the scrambled control. * Statistically significant at p-value < 0.05.

As an orthogonal approach, we modulated HepG2-CD81 cells with small molecule inhibitors targeting positive regulators identified in the screen (Fig. 3B) (Supplementary Table 1). IC50 values for each small molecule inhibitor were obtained in uninfected HepG2-CD81 cells using Live/ Dead staining (Fig. S2D). We included eltanexer, an inhibitor of exportin-1 (XPO1), a putative negative regulator of infection (Than et al., 2020). To test the role of LRP4 in sporozoite entry of hepatocytes, we pretreated HepG2-CD81 cells with HPA-12, a ceramide transport inhibitor (Berkes et al., 2016) and an LDL-R blocking peptide, which blocks LRP4, significantly inhibited sporozoite entry. Thus, both genetic and peptide-mediated intervention of LRP4 inhibits sporozoite entry of hepatocytes (Fig. 3A and B).

We next performed an imaging-based secondary screen with the selected 15 putative regulators to assess the role of these hits on the longer-term LS development. This was intended to more closely mirror the experiment performed in the initial screen, although we used a later time point in order to characterize the full impact on LS development. Specifically, individual CRISPR-Cas9 knockout lines were infected with *P. yoelii* sporozoites and observed 48 hours post infection (hpi). Several of the knockout lines exhibited substantially altered LS burden (Fig. 3C). The number of LS parasites was significantly increased in CENPJ (centromere protein J) and KDELC1 (Lys-Asp-Glu-Leu containing 1) disrupted lines, illustrating that each of these factors is indeed a negative regulator of infection. In contrast, knockout of CISD1 (CDGSH iron sulfur domain 1), COL4A3BP (collagen type IV alpha-3-binding protein), IREB (iron-responsive element-binding protein) and LRP4 significantly reduced the number of LS parasites 48 hpi (Fig. 3C). We next tested whether LS infection could be perturbed by targeting these factors with pharmacological inhibitors. Consistent with the genetic experiments, small molecule inhibitors (Supplementary table 1) that target CISD1, COL4A3BP, IREB, and LRP4, significantly reduced the number of LS parasites observed after 48 h of infection (Fig. 3D), further supporting the notion that these factors are positive regulators of LS infection.

We next asked if any of the screen hits altered the growth of LS parasites. Interestingly, knock out of VPS51 (Ang2) did not significantly alter parasite load but instead, the size of the parasite was significantly smaller (Fig. 3E). HepG2-CD81 cells expressing sgRNAs directed against COL4A3BP and LRP4, which both reduced the number of LS parasites (Fig. 3C), and the size of LS parasites (Fig. 3E). In contrast, depletion of CENPJ increased both the size and the number of LS parasites. Knockout of other putative regulators did not result in altered parasite size (Fig. 3E). Interestingly, while our screen was only set up to identify factors that altered infection rate, not LS growth, it is possible that some slow-growing parasites may have not reached the threshold of GFP levels to be included in the “infected” gate. Taken together, these studies identified several host factors influencing parasite entry, growth and development (Fig. 3F) and illustrate the utility of genome-scale functional screening for the discovery of host factors that regulate *Plasmodium* LS infection.

### γ-tubulin foci sequester at parasite periphery

Among several new regulators of infection identified from the screen, we choose to investigate CENPJ, one of the MT cytoskeleton organizing proteins. Depletion of CENPJ resulted in significant increase in parasite load (Fig. 3C) and growth (Fig. 3E). CENPJ is a conserved, ubiquitously expressed centrosomal protein with a key role in centriole organization and biogenesis (Cho et al., 2006; Ganem et al., 2009; Kohlmaier et al., 2009). The centrosome is a major microtubule organizing center (MTOC) (Hung et al., 2000). CENPJ depletion impairs centriole assembly, resulting in fragmented MTOCs and non-radial MT cytoskeleton organization (Cho et al., 2006; Ganem et al., 2009; Kohlmaier et al., 2009). To characterize the role of CENPJ in parasite development, we assessed the localization of γ-tubulin with γ-TuRC (γ-tubulin ring complex), a core functional unit of the MTOC (Wiese and Zheng, 2000). We infected HepG2-CD81 cells with *P. yoelii* sporozoites and allowed infection to proceed for 48 h. Cells were stained with anti-UIS4 (upregulated in infectious sporozoites gene 4) and γ-tubulin. In uninfected cells, γ-tubulin foci were localized primarily near nuclear periphery (88%) (Fig. 4A). Strikingly, in infected cells, a majority of γ-tubulin foci (~60%) were found in the cytoplasm associated with the parasitophorous vacuolar membrane (PVM) (Fig. 4A).

**Figure 4.**
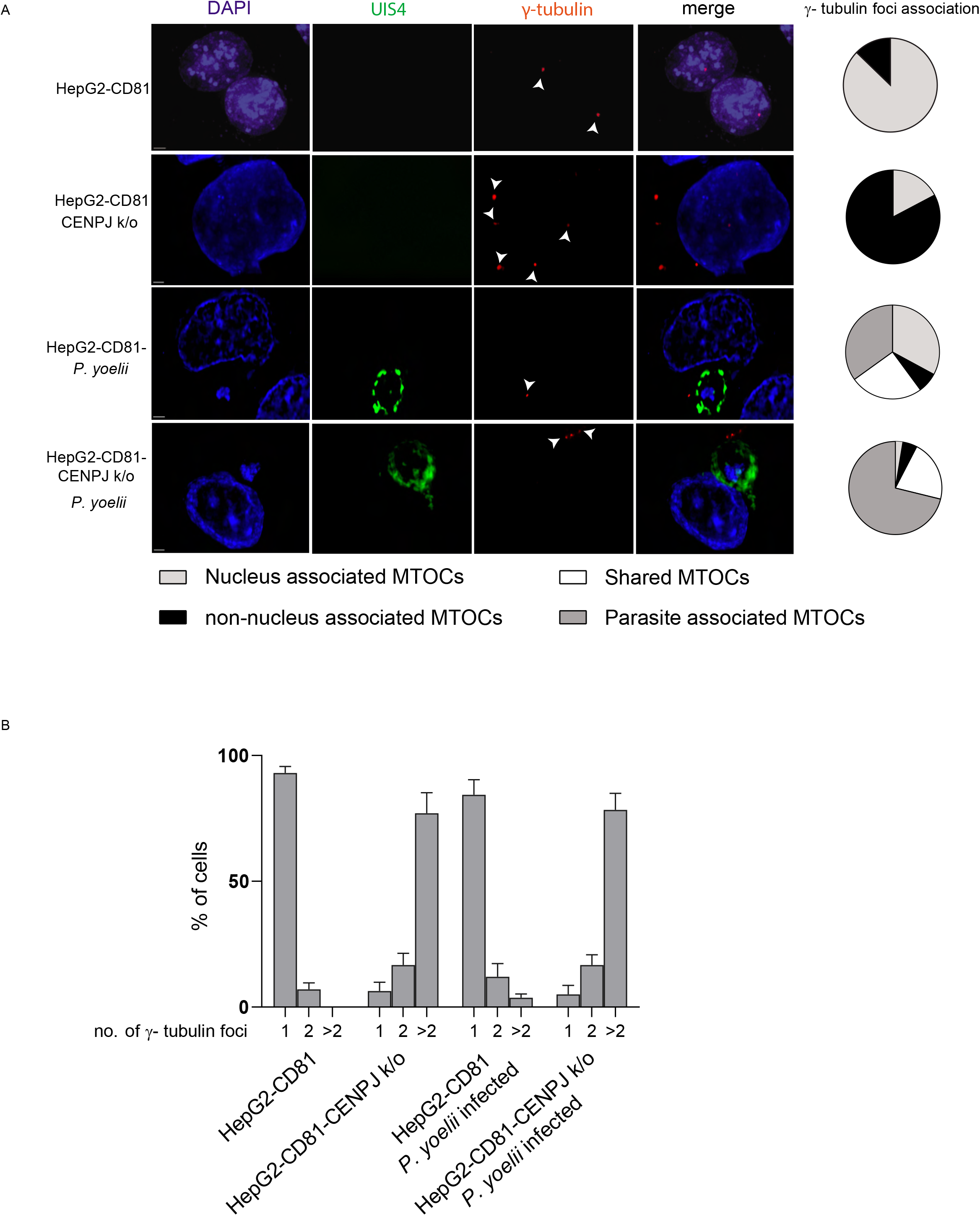
γ-tubulin foci sequester at parasite periphery. **(A)** HepG2-CD81 and HepG2-CD81-sgRNA-CENPJ cells were infected with *P. yoelii* sporozoites for 48 h. Cells were fixed and stained with antibodies against UIS4-Alexa 647 conjugate (pseudocolored green) and γ-tubulin to visualize the parasite PVM and the γ-tubulin foci, respectively. White arrow heads points to γ-tubulin foci. A pie chart showing localization of γ-tubulin in the cell during different conditions. Images are maximum intensity projections from 20-25 z-slices. Images are a representative of 3 independent experiments with n ≥ 100 cells *I* condition. Bar = 2 μm **(B)** A bar graph representing percentage of cells containing multiple γ-tubulin foci. Data shown represent 3 independent experiments with n ≥ 100 infected cells.

Next, we evaluated the functional role of CENPJ in regulating the LS parasite. In uninfected CENPJ depleted cells, we observed increased cytoplasmic localization (~80%) (Fig. 4A) and multiple γ-tubulin foci (Fig. 4B). Infection in CENPJ knockout cells resulted in an increase in γ-tubulin localization (~92%) to the PVM compared to infected control cells (~60%) (Fig. 4A) and an increase in LS infection (Fig.3C). Infection with LS parasites result in MTOC re-localization to the PVM (Fig 4A and B); the absence of CENPJ further exacerbated the non-centrosome MTOC organization close to PVM that supports LS development (Fig 4A and B).

### Golgi serves as a non-centrosomal MTOC (ncMTOC) in *P. yoelii* infected cells

Canonically, microtubule (MT) arrays nucleate from MTOCs and radiates towards cell periphery (Wiese and Zheng, 2000). To understand whether the γ-tubulin sequestration resulted in dynamic reorganization of MTs around the parasite, we infected rfp-α-tubulin transfected HepG2-CD81 cells with *P. yoelii* sporozoites, and allowed the infection to procced for 48 h. After 46 h, nocodazole was added. After an additional 2h (48h post-infection), cells were washed, and incubated with nocodazole-free media for 45 sec to allow the nucleation of MT. Cells were stained with γ-tubulin and UIS4. In uninfected cells, microtubules nucleated from γ-tubulin foci at the host nucleus (Fig. 5A). In the infected cells, MT nucleated from γ-tubulin foci localized adjacent to the PVM (Fig. 5A). This suggest that, during infection, MTOCs reorganizes the host microtubule network around the developing LS parasite.

**Figure 5.**
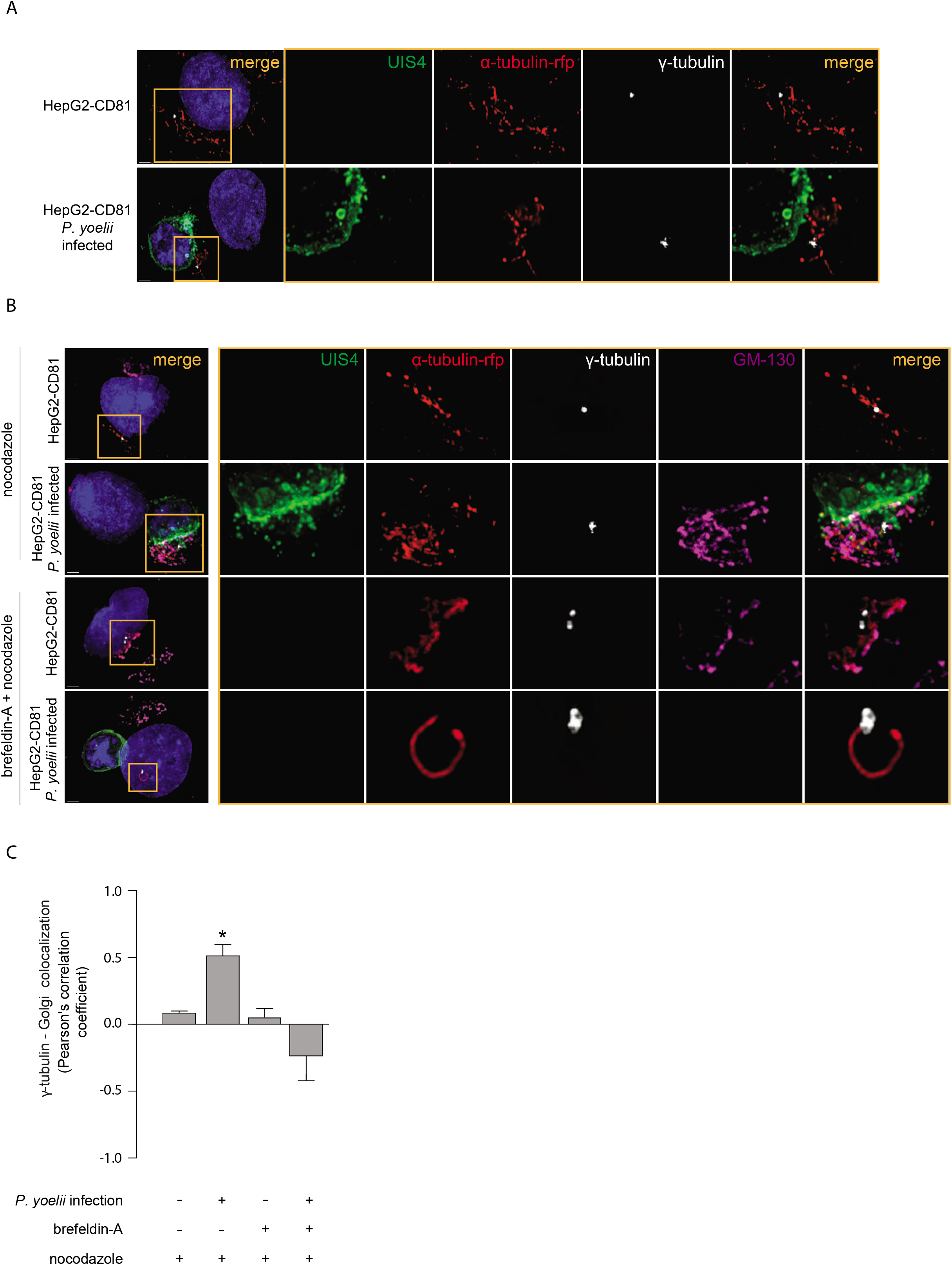
Golgi acts as ncMTOC in *P. yoelii* infected cells. **(A)** HepG2-CD81-rfp-tubulin cells were infected with *P. yoelii* sporozoites for 48 h. After 46 h, nocodazole was added. After an additional 2h (48h post-infection), cells were washed, and incubated for 45 sec to allow the nucleation of MT. Cells were stained with γ-tubulin and UIS4. Cells were fixed and stained with antibodies against UIS4-HepG2-CD81-rfp-tubulin cells were infected with *P. yoelii* sporozoites for 48 h; after 24h, cells were treated with brefeldin-A. 647 conjugate (pseudo colored-green) and γ-tubulin (white) to visualize the parasite PVM and the microtubule organizing center (MTOC), respectively. **(B)** After an additional 22 hours (46h post-infection), nocodazole was added to the cells. After an additional 2h, cells were washed, then incubated with media alone for 45 sec to allow MT nucleation. Cells were fixed and stained with antibodies against GM130, UIS4 and γ-tubulin to visualize the parasite PVM and the microtubule organizing center (MTOC), respectively. Images are representative of 3 independent experiments with n ≥ 100 infected and control cells. Bar = 2 μm. **(C)** Intensity based colocalization was performed on at least 75 cells per condition and Pearson’s correlation coefficients were calculated. The bar graph depicts mean ± SD of three independent experiments. The bar graph represents Pearson’s correlation coefficient calculated from **(B).**

Several studies have demonstrated that, in the absence of centrosome organizing proteins, Golgi outposts act as a non-centrosomal MTOCs (ncMTOCs) that function as MT nucleation sites by recruiting γ-tubulin foci ((Grimaldi et al., 2013) reviewed in (Zhu and Kaverina, 2013)). To test whether the cytoplasm localized γ-tubulin foci are regulated by Golgi outposts, we infected rfp-α-tubulin expressing HepG2-CD81 cells with *P. yoelii* sporozoites. We allowed the infection to procced for 48 h. As above, nocodazole was added to the cells after 46 hours. Cells were washed and incubated for an additional 45 sec to allow MT nucleation. Cells were fixed then stained with antibodies against the Golgi peripheral cytoplasmic membrane protein, Golgi membrane protein of 130 kDa; golgin subfamily A member 2 (GM130), γ-tubulin and UIS4. In uninfected cells, we observed nucleating MTs originating at γ-tubulin foci associated with the host nucleus. In *P. yoelii* infected cells, we primarily observed MTs nucleating from γ-tubulin foci in proximity to GM130 staining (Fig. 5B and C).

We next asked if an intact Golgi was required for PVM-associated MTOC formation. To do this, we utilized the small molecule brefeldin A, which reversibly disrupts and fragments the Golgi, blocking assembly and transport of secretory vesicles (Sciaky et al., 1997). We infected rfp-α-tubulin expressing HepG2-CD81 cells with *P. yoelii* sporozoites; after 24h cells were treated with brefeldin-A. Twenty two hours later (46h post-infection), nocodazole was added to the cells. After an additional 2h (48h postinfection), cells were washed, then incubated with media alone for 45 sec to allow MT nucleation. Cells were stained with antibodies against GM130, γ-tubulin and UIS4. In contrast to infected cells with intact golgi, following brefeldin-A treatment, we observed few γ-tubulin foci at the PVM and instead MT nucleation originating from at the nuclear periphery (Fig. 5B). These results are consistent with the hypothesis that the parasite utilizes Golgi-associated ncMTOC formation to initiate MT reorganization around PVM.

### Host Golgi and intracellular vesicles interact with *Plasmodium* liver stage

To better understand the role the Golgi plays in regulating ncMTOC formation, we studied Golgi-PVM interaction during infection. HepG2-CD81 cells were infected with *P. yoelii* sporozoites and stained cells with antibodies against the Golgi peripheral cytoplasmic membrane protein, GM130 (Golgi membrane protein of 130 kDa; golgin subfamily A member 2). We observed UIS4-positive membrane co-localized with Golgi stacks in nearly three quarters of the infected cells at 48 hpi (Fig. 6A). Consistent with other reports (De Niz et al., 2020; Raphemot et al., 2019), we also observed Golgi localized to the PVM (Fig. 6A). Co-localization between the PVM and Golgi was reduced following brefeldin-A treatment (Fig. 6A and C). These results are consistent with a model where an association between PVM-Golgi induces ncMTOC formation and reorganizes the MT network.

**Figure 6.**
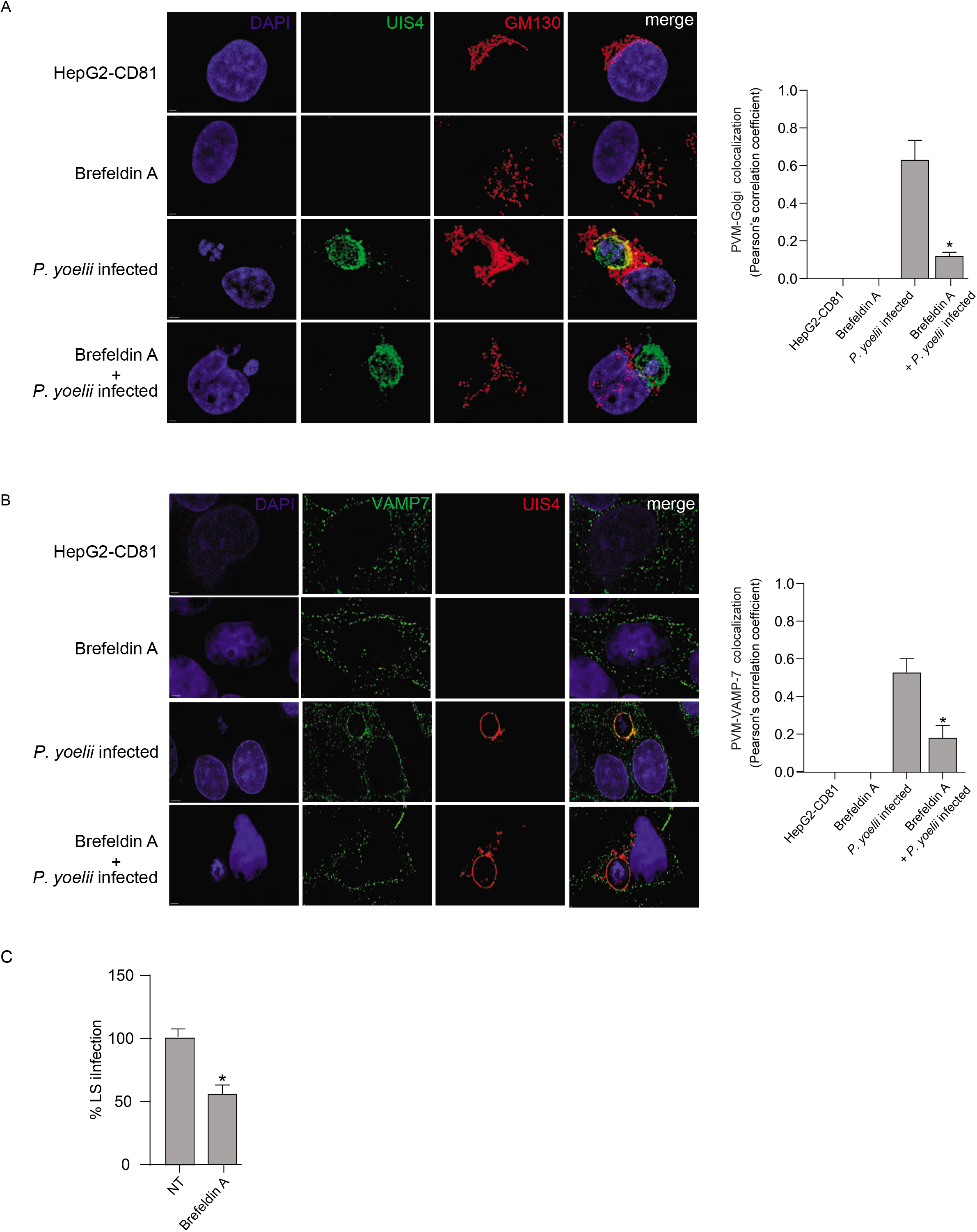
Host Golgi and intracellular vesicles interacts with *Plasmodium.* **(A)** HepG2-CD81 cells were infected with *P. yoelii* sporozoites for 48 h. Cells were treated with brefeldin-A following 24 h of infection. Cells were fixed and stained with Golgi GM130 and UIS4 to visualize host Golgi and the parasite PVM, respectively. A representative maximum intensity projection of is shown. **(B)** HepG2-CD81 cells were infected with *P. yoelii* sporozoites for 48 h. Cells were treated with brefeldin-A following 24 h of infection. Cells were fixed and stained with VAMP7 and UIS4 to visualize host intracellular vesicles and the parasite PVM, respectively. A representative maximum intensity projection of is shown for **(A)** and in-focus single z-image is shown for **(B)**. Images in **(A)** and **(B)** are representative of 3 independent experiments with n ≥ 100 infected and control cells. Bar = 2 μm. Intensity based colocalization was performed on at least 75 cells per condition and Pearson’s correlation coefficients were calculated. The bar graph depicts mean ± SD of three independent experiments. **(C)** The bar graph depicts the infection rate quantified from **(A)**. n=3; mean ± SD

We hypothesized that the reorganization of MT network by parasite serves to redirect vesicle traffic to the PVM and facilitate LS survival. This hypothesis is based on two observations: (1) that the Golgi reorients the MTOC to the parasite periphery and (2) that GO terms associated with vesicular trafficking and Golgi and ER stress were significantly enriched as putative regulators of infection in screen (Fig. 2B). We infected HepG2-CD81 cells with *P. yoelii* sporozoites and allowed infection to proceed for 24 or 36 h. Cells were stained with anti-UIS4 and anti-VAMP7 (to visualize intracellular vesicles) antibodies to visualize the PVM and host intracellular vesicles, respectively. We observed colocalization between intracellular vesicles and the PVM (Fig. 6B and C), consistent with other reports (De Niz et al., 2020; Raphemot et al., 2019). The colocalization of intracellular vesicles with the PVM was greatly reduced after brefeldin-A treatment (Fig. 6B and C), suggesting that Golgi integrity (and, likely, subsequent MTOC formation) is important for host vesicle trafficking to PVM. Together, these data suggest the parasite mediates Golgi-associated ncMTOC formation resulting in dynamic reorganization of MTs that redirects vesicular traffic to the PVM, promoting LS development.

## Discussion

For decades, dogma suggested that the elimination of anything short of 100% of LS parasites would result in little to no benefit in the effort towards malaria eradication. New evidence suggests this is not the case. Mathematical models suggest elimination of even a portion of hypnozoites could dramatically reduce *P. vivax* prevalence (White et al., 2018). Recently, it was demonstrated that targeting host aquaporin 3 leads to the elimination of *P. vivax* hypnozoites from field isolates (Posfai et al., 2020), suggesting that host targeted interventions may provide an opportunity to tackle even the Achilles heel of malaria control efforts. Another host factor known to be critical for LS infection, the tumor suppressor p53 (Douglass et al., 2015; Kain et al., 2020; Kaushansky et al., 2013), has been associated with lower severity of infection in Malian children (Tran et al., 2019). Additionally, host targeted interventions can induce at least partial immunity to subsequent challenge (Ebert et al., 2020). This supports the investigation into the use of host targeted drugs in casual prophylaxis strategies, provided the drugs have suitable toxicity profiles (reviewed in (Glennon et al., 2018)). Because not all host targets are suitable drug targets, a broad and comprehensive picture of factors that regulate LS malaria infection is needed.

Previous forward-genetic screens have identified host factors involved in *Plasmodium* infection (Prudencio et al., 2008; Raphemot et al., 2019; Rodrigues et al., 2008).These screens have exhibited very little overlap in identified factors (Fig. S1A, B & C), presumably in part because each screen prioritized identifying a small, but bona fide list of “hits,” and suffered a high false negative rate as a result. This has led to many key discoveries into interactions between the malaria parasite and its host hepatocyte but has fallen short of providing a systematic view of the fundamental biological properties that regulate the development and survival of the LS parasite. Like the earlier screens, the CRISPR-Cas9 screen we report here does not exhibit substantial overlap with previous screens when individual gene hits are evaluated, suggesting that additional analysis is still needed to comprehensively assess factors that regulate infection. Yet, when we evaluate whether hits from our screen are present in similar pathways to those observed in other screens, the overlap is substantial (Fig. S1D & E). Thus, while we may have, as a field, identified many central regulatory biological functions that control LS development, we have yet to saturate our understanding of the molecular players that mediate these biological necessities. Together, the CRISPR-Cas9 screen we present here, along with the previously reported siRNA screens, represent a key resource for the field moving forward, and we anticipate that merging findings from these experiments (Supplemental File 1) will provide many additional hypotheses to probe. One limitation of this work is that, since it is likely that at least a subset of canonical signaling pathways are rewired in the course of infection (Glennon et al., 2019), pathway analysis, which is based primarily on canonical signaling networks, is unlikely to comprehensively describe the an entirely accurate topology of the signaling relationships that mediate the complex host-parasite interface. Developing tools to reconstruct signaling relationships, within the context of malaria infection, is a critical area for future investigation.

Multiple screening efforts and other reports (Lopes da Silva et al., 2012; Niklaus et al., 2019; Petersen et al., 2017; Prudencio et al., 2008; Raphemot et al., 2019; Real et al., 2018; Rodrigues et al., 2008; Vijayan et al., 2019) have pointed to the critical role of vesicular transport, including retrograde trafficking (Raphemot et al., 2019), throughout *Plasmodium* LS infection. Here, we observe an association of Golgi and post-Golgi anterograde vesicles with the parasite, suggesting the parasite sequesters both anterograde and retrograde trafficking of host derived vesicles. Our data are consistent with the model that these vesicles are trafficked via MT to the parasite periphery as a result of ncMTOCs localized to the parasite periphery, assembled by the closely associated Golgi. This MTOC localization and subsequent trafficking provides a functional benefit to the parasite, as knockdown of CENPJ, which inhibits centrosomal MTOC formation and further exacerbates the localization of ncMTOCs to the parasite periphery, increases LS parasite infection. Interestingly, several reports previously identified the fusion of host derived vesicles such as late endosomes (Lopes da Silva et al., 2012), lysosomes (Niklaus et al., 2019), autophagosomes (Real et al., 2018) and retrograde vesicles from Golgi with the parasite (Raphemot et al., 2019) to enable development. Our data suggest that this fusion is facilitated by a Golgi-mediated ncMTOC assembly at the PVM and the resultant vesicular trafficking to the parasite periphery. Similar reorganization of the host MT network to facilitate the hijacking of host vesicles has been reported in *Toxoplasma gondii* (Coppens et al., 2006), suggesting that this may be a conserved mechanism by which Apicomplexan parasites exploit for nutrient uptake and survival.

### Significance

New strategies to combat malaria in the field are desperately needed. The causative agent of malaria, the obligate intracellular parasite *Plasmodium,* relies heavily on its mammalian host to survive and develop. The identification host regulators of infection provide an opportunity combat parasites, particularly during their initial liver stage of infection. Here, we perform a genome wide forward genetic screen to identify host factors that regulate *Plasmodium* liver stage infection. We demonstrate a mechanism by which the parasite remodels the host cytoskeleton to redirect host vesicular traffic to the parasite and facilitate its development. Our work implicates diverse host processes in liver stage *Plasmodium* development, which may be leveraged to develop pharmacological agents to fight malaria.

## Materials and Methods

### Cell lines, plasmids and antibodies

HepG2-CD81 cells (Silvie et al., 2006) were maintained in DMEM-Complete Medium (Dulbecco’s modified eagle medium (Cellgro, Manassas, VA), supplemented with 10% v/v FBS (Sigma-Aldrich, St. Louis, MO), 10000 IU/ml penicillin, 100 mg/ml streptomycin (Cellgro), 2.5 mg/ml fungizone (HyClone/Thermo Fisher, Waltham, MA) and 4 mM L-Glutamine (Cellgro). Cells were split 2-3 times weekly. All the plasmids for Genome-wide CRISPR library generation (GeCKOv2) and individual gene knockouts were procured from Addgene (MA, USA). CellLight Tubulin-RFP / GFP, BacMam 2.0 ready transfection mix were purchased from Thermoscientific (Missouri, USA). The following antibodies were used throughout this study: From Cell Signaling Technologies: acetylated alpha tubulin (5335S); Thermoscientific: gamma tubulin (PA1-28042), abcam: VAMP7 (ab36195) and Novus biologicals: GM130 (NBP2-53420) LSBio: UIS4 (LS-C204260-400).

### Mosquito rearing and sporozoite production

For *P. yoelii* sporozoite production, female 6–8-week-old Swiss Webster mice (Harlan, Indianapolis, IN) were injected with blood stage *P. yoelii* (17XNL) parasites to begin the growth cycle. Animal handling was conducted according to the Institutional Animal Care and Use Committee-approved protocols. Briefly, *Anopheles stephensi* mosquitoes were allowed to feed on infected mice after gametocyte exflagellation was observed. Salivary gland sporozoites were isolated using a standard protocol at day 14 or 15 post-blood meal. The sporozoites were activated with 20% v/v FBS and pelleted by centrifugation at 1,000 × *g* to salivary gland detritus. Sporozoites were further enriched by a second centrifugation at 15,000 × *g* for 4 min at 4 °C, before resuspension in a desired volume of complete medium.

### Pooled genome-wide CRISPR screen

To perform the whole-genome CRISPR screen, HepG2-CD81 cells were transduced with lentivirus containing the GeCKOv2 pooled sgRNA library of 123,642 sgRNAs targeting 19,031 protein-coding genes (~6 sgRNAs/gene), 1,864 microRNAs (4 sgRNA/microRNA) and 1,000 negative controls (2 sgRNA/control), and selected in puromycin for 5-7 days. On day 12-14 post-transduction, 40 million puromycin-resistant cells were infected with GFP tagged-*P. yoelii* at a MOI of 0.3. After 24 h of infection, cells were sorted as infected and uninfected by FACS into different bins based on GFP signal. A non-treated, noninfected control was also collected for each experiment to assess library representation. The experiment was performed four independent times. Genomic DNA from each sample was isolated using QIAamp DNA mini kit (Qiagen, Hilden, Germany).

### Next-generation sequencing

Libraries were generated using a 2-step PCR according to previously published protocol (Sanjana et al., 2014). Briefly, an initial PCR was performed using AccuPrime Pfx Supermix (Invitrogen, Waltham, MA, USA) with lentiCRISPRv2 adaptor primers to amplify the sgRNA region and add priming sites for Illumina indexing. Amplicons were purified using FlashGels (Lonza, Allendale, NJ, USA) and purified PCR products were used as templates for subsequent PCR amplification. Sufficient PCR reactions were performed to maintain library coverage. Next, a second PCR was performed in order to add Illumina P5 and P7 index sequences, as well as barcodes for multiplexing, and samples were repurified. Purified libraries were quantified using the KAPA library quantification kit (Kapa Biosystems, Wilmington, MA, USA) as per manufacturer’s instructions performed on an Applied Biosystems 7500 Fast real-time PCR machine (Applied Biosystems, Foster City, CA, USA). Samples were sequenced on a MiSeq (Illumina) using the manufacturer’s protocol with addition of Illumina PhiX control (Illumina, San Diego, CA, USA) to improve library diversity at a final concentration of 10% per library volume. After demultiplexing, FASTQ data files of 75bp single mate reads averaged 24.98 million raw reads per library.

### Differential abundance of guides and gene enrichment analysis

FASTQ files were aligned to the GeCKOv2 pooled sgRNA library of 123, 642 sgRNA DNA sequences by Bowtie2 (version 2.2.8) using local alignment policy command line arguments “ --local -L 12 -N 0 -D 15 -i C,1,0 --gbar 8 --rdg 10,3 --rfg 10,3”. This yielded on average 20.41 million aligned reads to guides per library. Read counts per guide were converted to relative expression abundance as Reads Per Million (RPM). A guide was called detected in a screen if the before infection condition was at least 0.1 RPM. Undetected guides (RPM below 0.1) were excluded from further calculations. 16,629 out of 19,031 (87.38%) genes targeted by 3 or more guides/ sgRNA were detected in least 3 experiments. Fold change with respect to ‘before infection’ was calculated by dividing RPM in ‘infected’ or ‘bystanders’ conditions by RPM in ‘before infection’ condition. The differential abundance of a guide is represented as the log_2_ ratio of fold change in ‘infected’ condition divided by the fold change in ‘bystanders’ condition. If less than two screens call a guide detected (RPM >= 0.2), a log_2_FC of 0 and p-value of 1 are reported for this guide. Otherwise, the final log_2_FC of the guide is the arithmetic mean of the log_2_ ratios from each detected screen, and the final p-value of the guide is calculated by one sample t-test that the log_2_ ratios of the detected guides was not zero. The GeCKO library contains 6 independent guides for each protein-coding gene. The log_2_FC and p-value at the gene level is calculated from log_2_FC and p-value of its 6 guides. The log_2_FC of a gene is equal to the log_2_FC of the guide with the lowest (best) p-value. The corrected p-value of a guide is set to 1 if the sign of its log_2_FC is opposite to the log_2_FC of the gene. Then the p-value of a gene is calculated as the product of corrected p-values from all guides not excluded from calculations.

Gene set enrichment analysis on all genes with positive/negative log_2_FC was performed based on major knowledgebases including HUGO Gene Nomenclature Committee (HGNC), Gene Ontology (GO), Kyoto Encyclopedia of Genes and Genomes (KEGG) and Reactome. The top 200 significantly enriched gene sets associated with all genes of negative or positive log_2_FC were identified. Genes that are both statistically significant (p-value < 0.05) and differentially abundant (log_2_FC < −6.0 or > 6.0) were considered significantly represented in *P. yoelii* infection. Go terms were clustered into higher order hierarchy using ClueGO plug-in (version 2.3.3), implemented in Cytoscape v3.4.0.

### Generation of individual hits using gene specific CRISPR sgRNA

GFP tagged vectors for the 15 hits were obtained from ABM Good (Richmond, British Columbia, Canada). Non-replicating lentiviral stocks were generated by transfection of HEK293-FT cells. 4 × 10^6^ HEK293-FT cells were plated on poly-L-lysine coated dishes to achieve 70-80% confluency at time of transfection. Approximately 24 h after plating, transfection mixtures were prepared by mixing 20 μl Polyethyleneimine MAX (Polysciences Inc, Warrington, PA) prepared at 1 mg/ml, together with 4.75 μg of sgRNA construct or a scramble control, along with 3^rd^ generation lentiviral packaging mix from ABM Good, according to manufacturer’s protocol. After incubating for 10 min at room temp in DMEM, transfection complexes were added dropwise to cells. After overnight incubation, cells were washed to remove transfection mixtures and were fed with 10 ml fresh media. Lentivirus-containing supernatant was harvested 36 hours later, passed through 0.45 μm syringe filters, and either used immediately for transduction or stored at −80 °C. To disrupt candidate genes, HepG2-CD81 cells were transduced with lentiviral supernatants in 6-well plates at a cell density of 1 × 10^6^ per well. At time of plating, cells were transduced with 1 ml of supernatant in the presence of 0.5 μg/ml polybrene (Sigma Aldrich St. Louis, MO). In order to select for cells with stable integration of shRNA transgenes, supernatant was replaced with complete media with the addition of 2 μg/ml puromycin 24 h post-transduction, and cells were selected for at least 5 days prior to experiments. For analysis of experiments with the knockout cells, only the GFP-positive cells have been considered.

### Infection assay

5 × 10^5^ HepG2-CD81 wild type cells or knockdown cells were seeded in each well of a 24-well plate (Corning) and infected with *P. yoelii* sporozoites at an MOI of 0.3 for 90 min and the infection was either stopped, or media was replaced and the infection was allowed to progress for 48 h. For drug treatment experiments, cells were either pretreated for 24 h and washed before infection or post treated following 24 h of infection as mentioned and the infection was allowed to proceed for 48 h.

### Flow cytometry

Cells were detached with accutase (Life technologies) and fixed with Cytoperm/Cytofix (BD Biosciences). Cells were blocked with Perm/Wash (BD Biosciences) + 2% (w/v) BSA for one hour at room temperature then stained overnight at 4 °C with Alexa Fluor −488 or −647 conjugated circumsporozoite (CSP) antibody. The cells were then washed and resuspended in PBS supplemented with 5 mM EDTA. Infection rates were measured by flow cytometry on an LSRII (Becton-Dickinson) and analyzed with FlowJo (Tree Star).

### Immunofluorescence

For imaging experiments, HepG2-CD81 wild type or knockout cells were plated in 8 well chamber slides (Labtek) and infected with *P. yoelii*sporozoites. Cells were fixed with 3.7% (v/v) paraformaldehyde (Sigma) at defined timepoints after infection (90 min or 48 h), permeabilized with Triton X-100, and stained with fluorescent tagged UIS-4 or other antibodies mentioned. Nuclei were stained with DAPI (Vectashield). Images were acquired with a 100× 1.4 NA objective (Olympus) on a DeltaVision Elite High-Resolution Microscope (GE Healthcare Life Sciences). The sides of each pixel represent 64.5 × 64.5 nm and z-stacks were acquired at 300 nm intervals. Approximately 20-30 slices were acquired per image stack. For deconvolution, the 3D data sets were processed to remove noise and reassign blur by an iterative Classic Maximum Likelihood Estimation widefield algorithm provided by Huygens Professional Software (Scientific Volume Imaging BV, The Netherlands). Images for processed with IMARIS Bitplane, image analysis software to quantify LS, perform colocalization analysis and remove outlier cells. For the high throughput secondary screen, cells were plated onto 96 well plate, infected and stained as explained above. Images were acquired using Keyence BZ-X800 automated microscope and infection rate were quantified using Imaris 9.5, image analysis software.

### Microtubule nucleation assay

For analysis of Golgi nucleation of microtubules, cells were treated with 20 nM nocodazole for 2 h at 37 °C and immediately incubated with extraction buffer (60 mM PIPES, 25 mM HEPES, 10mM EGTA, 2 mM MgCl2, 0.1% Tritón X-100, pH 6.9, supplement with 0.25 nM nocodazole and 0.25 nM taxol) for 45 s. Cells were then fixed with methanol and stained with antibody anti GM130 and antibody anti α-tubulin. For brefeldin-A treatment experiments, HepG2-CD81-rfp-tubulin cells were infected with *P. yoelii* sporozoites for 48 h; after 24h, cells were treated with brefeldin-A. 22 h after brefeldin-A treatment, cells were treated with 20 nM nocodazole for 2 h at 37 °C and immediately incubated with extraction buffer (60 mM PIPES, 25 mM HEPES, 10mM EGTA, 2 mM MgCl2, 0.1% Tritón X-100, pH 6.9, supplement with 0.25 nM nocodazole and 0.25 nM taxol) for 45 s. Cells were then fixed with methanol and stained with antibody mentioned. Images were acquired with a 100× 1.4 NA objective (Olympus) on a DeltaVision Elite High-Resolution Microscope (GE Healthcare Life Sciences) and analyzed as explained above.

### Meta-analysis of screens

Z-scores of positively and negatively represented genes in each screen were calculated separately. Meta ranking was performed by function metaRank() from the R package DuffyTools, (using arguments: mode=“percentile”, rank.average.FUN=mean, naDropPercent = 0.75). Positively represented genes with z-scores greater than the cutoff (1.0, 1.5, 2.0) and negatively represented genes with z-scores smaller than the cutoff (-1.0, −1.5, −2.0) were selected as hits for each screen. Hits of positively and negatively represented genes were combined for further pathway enrichment analysis. To compare datasets of uneven sizes, gene rank percentiles were assigned to positively and negatively represented genes in each screen separately. Genes were ranked by the average percentiles across all datasets where they were screened.

### Gene Ontology analysis on integrated forward genetic screens

Identified hit genes from all the four screens were uploaded in ious combinations in the ClueGO plug-in (version 2.3.3), implemented in Cytoscape v3.4.0 (http://cytoscape.org/) to generate gene ontology (GO) and pathway enrichment networks. Enriched functionally annotated groups were obtained with the following setting parameters: organism was set to Homo sapiens; the total gene set used in each of the screen were used as reference; the gene ontology terms were accessed from the following ontologies/pathways: Biological Process and Reactome Pathway database evidence code was restricted to ‘All_without_IEA’. The GO fusion option was also selected. The significance of each term was calculated with a two-sided hypergeometric test corrected with Benjamini-Hochberg correction for multiple testing. The kappa score was set to 0.5 and the GO tree levels were restricted at 6–16 (medium-detailed specificity). For GO term selection, a minimum of 3 genes and 3% coverage of the gene population was set. GO terms were grouped with an initial group size of 2 and 50% for group merge. The remaining parameters were set to defaults.

## Statistical analysis

GraphPad Prism 7 was used for all statistical analyses. Statistical details for each experiment can be found in the figure legends, including the number of technical and biological replicates performed.

## Acknowledgements

This research was funded by National Institutes of Health grants R01GM101183 (AK), R00AI111785 (AK), a W.M. Keck Foundation award (AK), U19AI100627 (AA) R01AI032972 (AA), U19AI111276 (JDA), and P41GM109824 (JDA). We thank the Seattle Children’s Research Institute vivarium staff for their work with mice and insectary staff for their work with mosquitoes.

---

**Figure S1.**
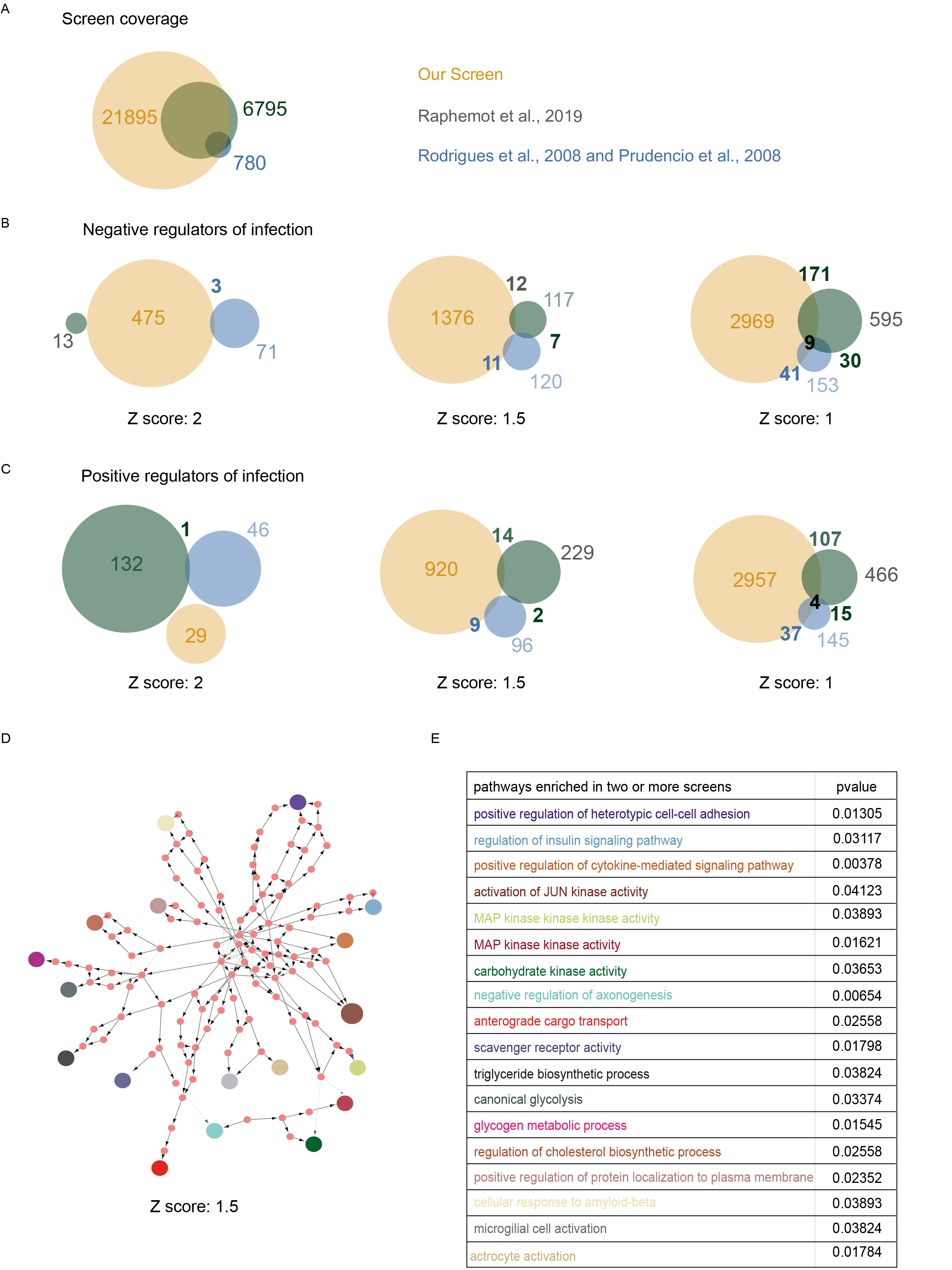
Meta-analysis of *Plasmodium* liver stage screens reveals areas for future investigation. **(A)** A Venn diagram showing gene-level coverage in three screening efforts. **(B)** A Venn diagram depicts gene-level overlap of the negative regulators of infection identified following meta-analysis (Supplementary file 1) on the gene hits from three screens compared with the z-score cut-off of 2, 1.5 and 1. **(C)** A Venn diagram showing overlap of positive regulators of infection identified following meta-analysis (Supplementary file 1) on the gene hits from the three screens compared with the z-score cut-off of 2, 1.5 and 1. **(D)** Network analysis on the enriched GO biological processes predicted from significantly enriched genes identified from at least two of three screens with the z-score cut-off of 1.5. Nodes represent biological processes, interactions between pathways. The gene and pathways identified from the analysis shared by the biological processes are represented solid arrows while the gene shared by the biological processes are represented as dashed lines. The significant biological processes identified from the network analysis **(D)** are color coded in **(E)** and tabulated with corresponding p values.

**Figure S2:**
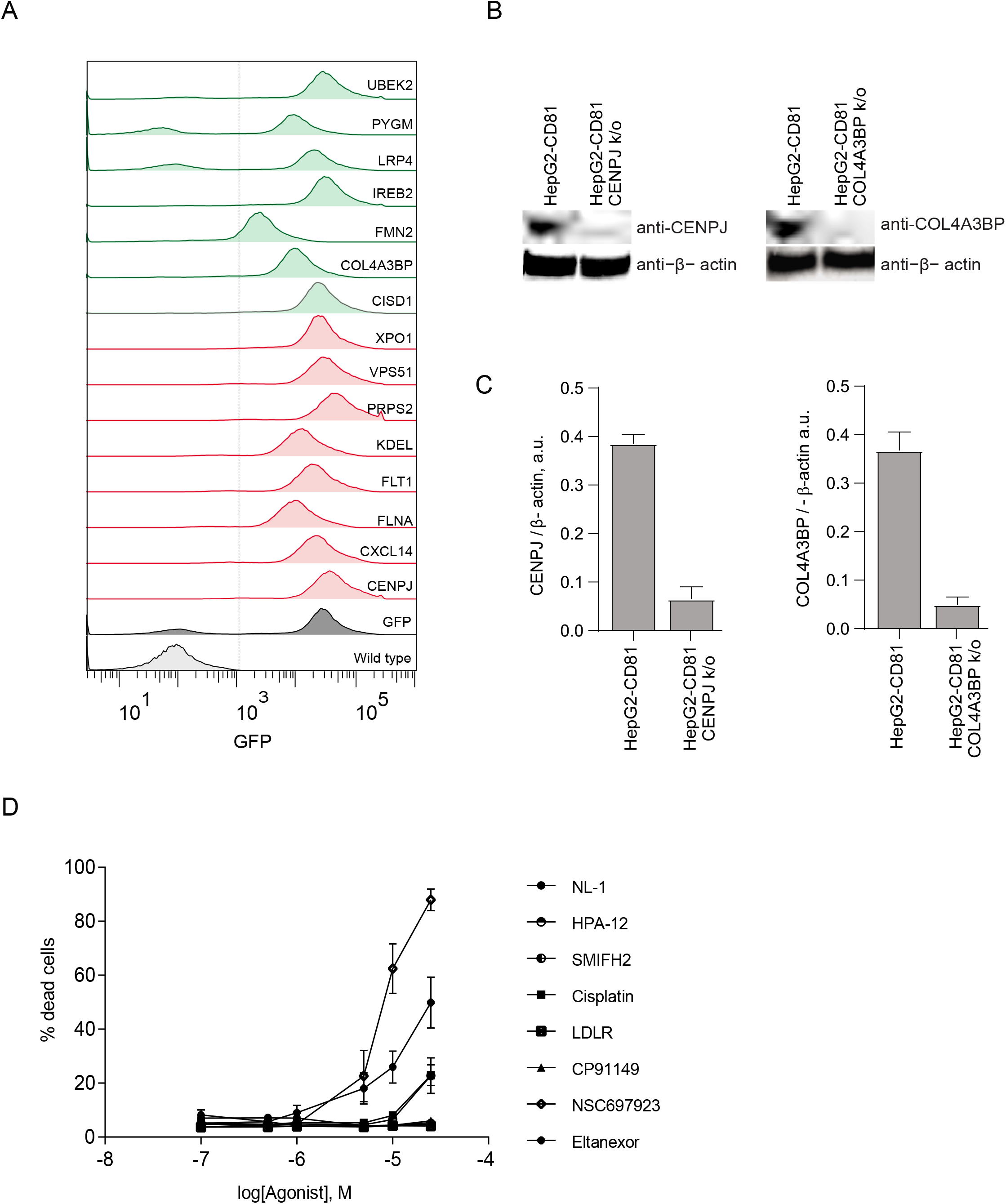
**(A)** HepG2-CD81 cells were transfected with CRISPR-Cas9 – GFP-sgRNA containing plasmids targeting the specified gene or scrambled control. A representative histogram shows levels of GFP incorporation post transfection confirming the successful integration of sgRNA into the transfected cell. **(B)** Western blot with anti-CENPJ and anti-COL4A3BP antibody showing the expression levels of CENPJ and COL4A3BP in scrambled and CENPJ knockout cells. **(C)** The bar graph represents the quantification of CENPJ and COL4A3BP expression in wildtype and knockout cells. n=3; mean ± SD. **(D)** Evaluation of cytotoxicity profile of small molecules used in the study. Data are presented as the mean cytotoxicity value ± standard deviation from one representative experiment of three independent experiments.

**Supplementary table 1:**
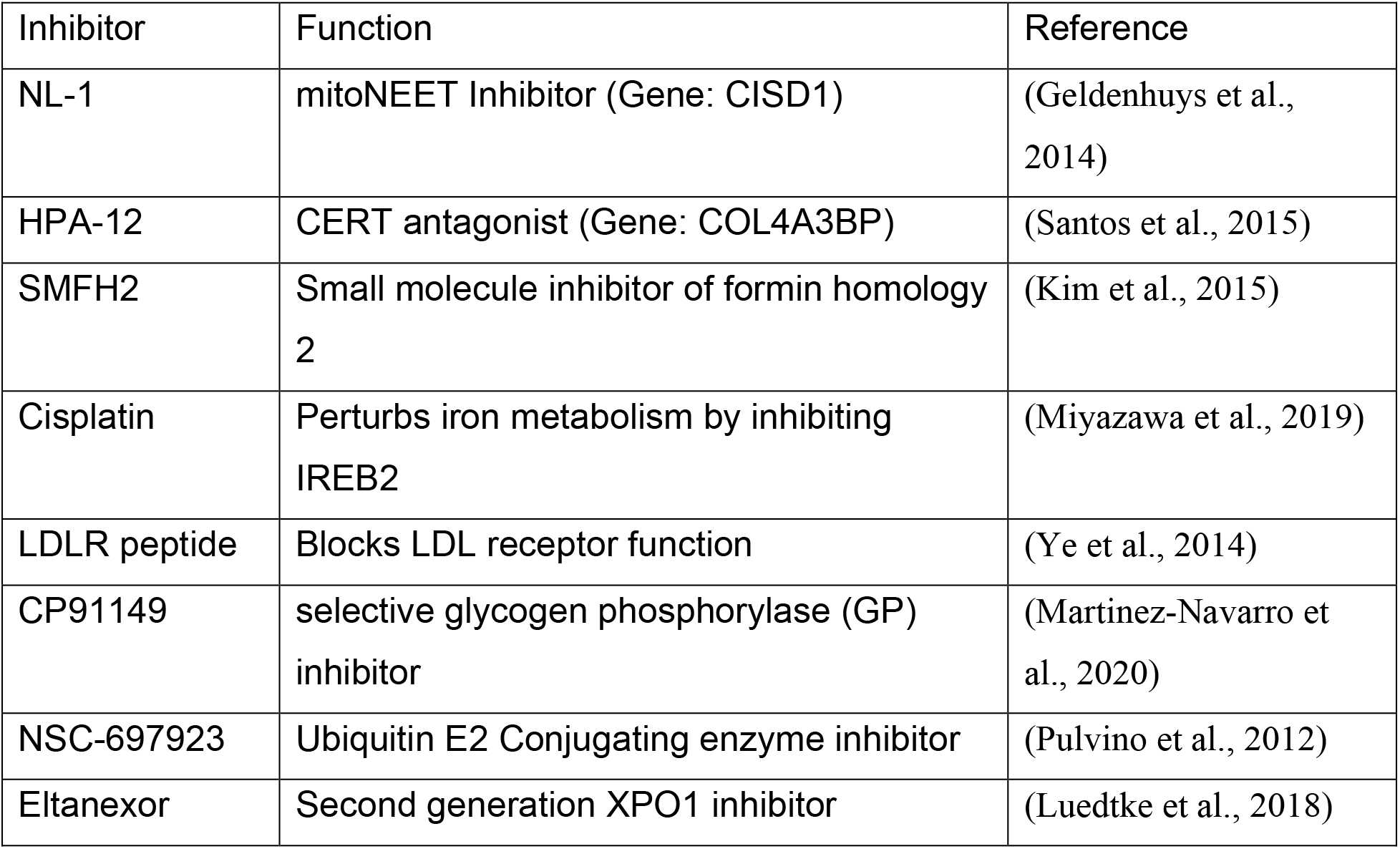
Inhibitors used in the study. Related to figure 3.

**Supplementary table 2:**
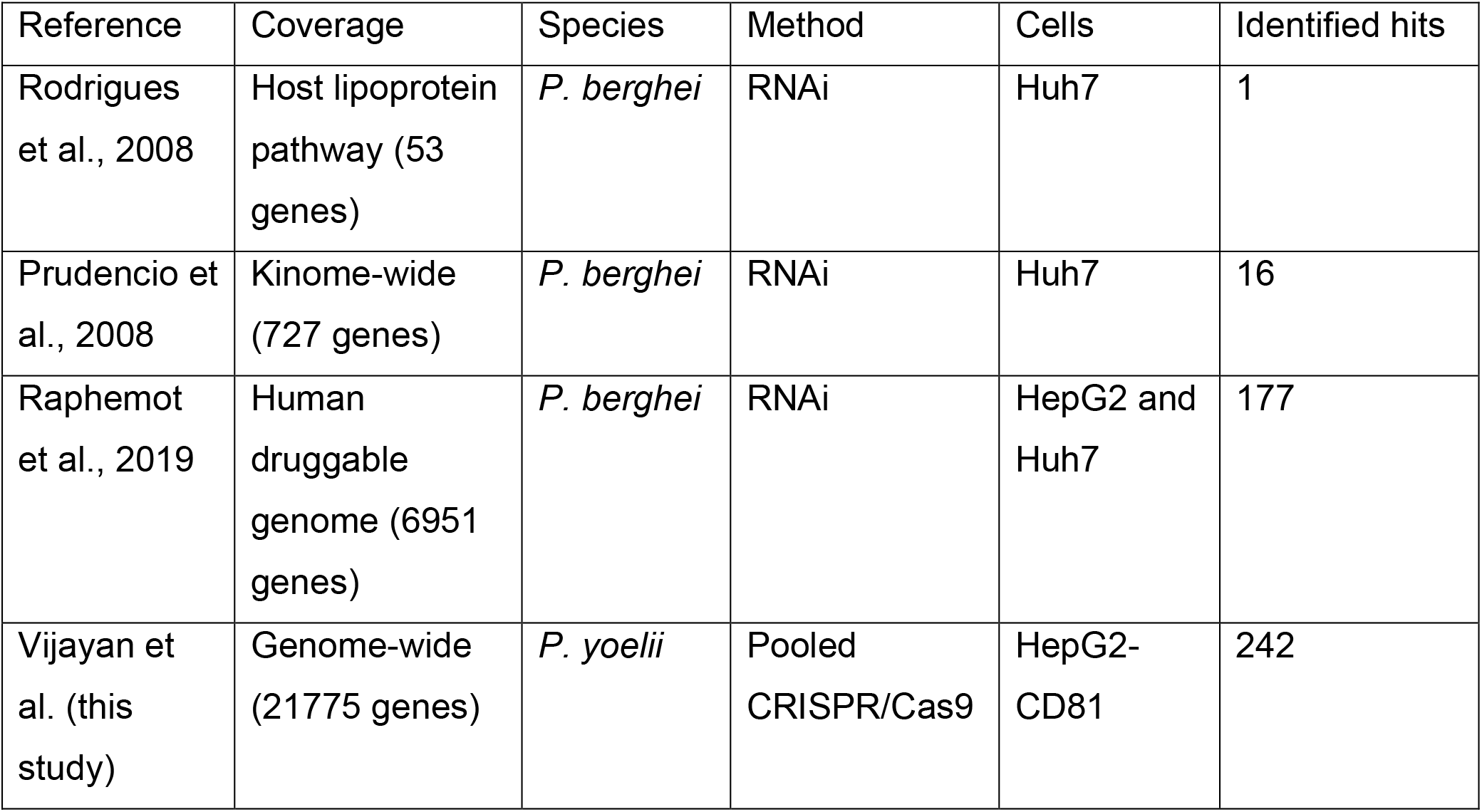
Comparison of our CRISPR/Cas9 screen with previous hostbased genetic intervention screens. Related to figure S1.

| Reference | Coverage | Species | Method | Cells | Identified hits |
| --- | --- | --- | --- | --- | --- |
| Rodrigues et al., 2008 | Host lipoprotein pathway (53 genes) | *P. berghei* | RNAi | Huh7 | 1 |
| Prudencio et al., 2008 | Kinome-wide (727 genes) | *P. berghei* | RNAi | Huh7 | 16 |
| Raphemot et al., 2019 | Human druggable genome (6951 genes) | *P. berghei* | RNAi | HepG2 and Huh7 | 177 |
| Vijayan et al. (this study) | Genome-wide (21775 genes) | *P. yoelii* | Pooled CRISPR/Cas9 | HepG2-CD81 | 242 |

## References

Alix, E., Mukherjee, S., and Roy, C.R. (2011). Subversion of membrane transport pathways by vacuolar pathogens. The Journal of cell biology 195, 943–952.

Asrat, S., de Jesus, D.A., Hempstead, A.D., Ramabhadran, V., and Isberg, R.R. (2014). Bacterial pathogen manipulation of host membrane trafficking. Annu Rev Cell Dev Biol 30, 79–109.

Berkes, D., Daich, A., Santos, C., Ballereau, S., and Genisson, Y. (2016). Chemistry and Biology of HPAs: A Family of Ceramide Trafficking Inhibitors. Chemistry 22, 17514–17525.

Bindea, G., Mlecnik, B., Hackl, H., Charoentong, P., Tosolini, M., Kirilovsky, A., Fridman, W.H., Pages, F., Trajanoski, Z., and Galon, J. (2009). ClueGO: a Cytoscape plug-in to decipher functionally grouped gene ontology and pathway annotation networks. Bioinformatics 25, 1091–1093.

Burda, P.C., Caldelari, R., and Heussler, V.T. (2017). Manipulation of the Host Cell Membrane during Plasmodium Liver Stage Egress. mBio 8.

Cho, J.H., Chang, C.J., Chen, C.Y., and Tang, T.K. (2006). Depletion of CPAP by RNAi disrupts centrosome integrity and induces multipolar spindles. Biochem Biophys Res Commun 339, 742–747.

Coppens, I., Dunn, J.D., Romano, J.D., Pypaert, M., Zhang, H., Boothroyd, J.C., and Joiner, K.A. (2006). Toxoplasma gondii sequesters lysosomes from mammalian hosts in the vacuolar space. Cell 125, 261–274.

De Niz, M., Kaiser, G., Zuber, B., Do Heo, W., Heussler, V.T., and Agop-Nersesian, C. (2020). Hijacking of the host cell Golgi by <em>Plasmodium</em> liver stage parasites. bioRxiv, 2020.2007.2024.220137.

Douglass, A.N., Kain, H.S., Abdullahi, M., Arang, N., Austin, L.S., Mikolajczak, S.A., Billman, Z.P., Hume, J.C.C., Murphy, S.C., Kappe, S.H.I., et al. (2015). Host-based Prophylaxis Successfully Targets Liver Stage Malaria Parasites. Mol Ther 23, 857–865.

Ebert, G., Lopaticki, S., O’Neill, M.T., Steel, R.W.J., Doerflinger, M., Rajasekaran, P., Yang, A.S.P., Erickson, S., Ioannidis, L., Arandjelovic, P., et al. (2020). Targeting the Extrinsic Pathway of Hepatocyte Apoptosis Promotes Clearance of Plasmodium Liver Infection. Cell Rep 30, 4343–4354 e4344.

Ganem, N.J., Godinho, S.A., and Pellman, D. (2009). A mechanism linking extra centrosomes to chromosomal instability. Nature 460, 278–282.

Glennon, E.K.K., Austin, L.S., Arang, N., Kain, H.S., Mast, F.D., Vijayan, K., Aitchison, J.D., Kappe, S.H.I., and Kaushansky, A. (2019). Alterations in Phosphorylation of Hepatocyte Ribosomal Protein S6 Control Plasmodium Liver Stage Infection. Cell Rep 26, 3391–3399 e3394.

Glennon, E.K.K., Dankwa, S., Smith, J.D., and Kaushansky, A. (2018). Opportunities for Host-targeted Therapies for Malaria. Trends Parasitol 34, 843–860.

Gomes-Santos, C.S., Itoe, M.A., Afonso, C., Henriques, R., Gardner, R., Sepulveda, N., Simoes, P.D., Raquel, H., Almeida, A.P., Moita, L.F., et al. (2012). Highly dynamic host actin reorganization around developing Plasmodium inside hepatocytes. PLoS One 7, e29408.

Grimaldi, A.D., Fomicheva, M., and Kaverina, I. (2013). Ice recovery assay for detection of Golgi-derived microtubules. Methods Cell Biol 118, 401–415.

Hale, V.L., Watermeyer, J.M., Hackett, F., Vizcay-Barrena, G., van Ooij, C., Thomas, J.A., Spink, M.C., Harkiolaki, M., Duke, E., Fleck, R.A., et al. (2017). Parasitophorous vacuole poration precedes its rupture and rapid host erythrocyte cytoskeleton collapse in Plasmodium falciparum egress. Proc Natl Acad Sci U S A 114, 3439–3444.

Hung, L.Y., Tang, C.J., and Tang, T.K. (2000). Protein 4.1 R-135 interacts with a novel centrosomal protein (CPAP) which is associated with the gamma-tubulin complex. Mol Cell Biol 20, 7813–7825.

Itoe, M.A., Sampaio, J.L., Cabal, G.G., Real, E., Zuzarte-Luis, V., March, S., Bhatia, S.N., Frischknecht, F., Thiele, C., Shevchenko, A., et al. (2014). Host cell phosphatidylcholine is a key mediator of malaria parasite survival during liver stage infection. Cell host & microbe 16, 778–786.

Kain, H.S., Glennon, E.K.K., Vijayan, K., Arang, N., Douglass, A.N., Fortin, C.L., Zuck, M., Lewis, A.J., Whiteside, S.L., Dudgeon, D.R., et al. (2020). Liver stage malaria infection is controlled by host regulators of lipid peroxidation. Cell Death Differ 27, 44–54.

Kaushansky, A., Ye, A.S., Austin, L.S., Mikolajczak, S.A., Vaughan, A.M., Camargo, N., Metzger, P.G., Douglass, A.N., MacBeath, G., and Kappe, S.H. (2013). Suppression of host p53 is critical for Plasmodium liver-stage infection. Cell Rep 3, 630–637.

Kohlmaier, G., Loncarek, J., Meng, X., McEwen, B.F., Mogensen, M.M., Spektor, A., Dynlacht, B.D., Khodjakov, A., and Gonczy, P. (2009). Overly long centrioles and defective cell division upon excess of the SAS-4-related protein CPAP. Curr Biol 19, 1012–1018.

Labaied, M., Jayabalasingham, B., Bano, N., Cha, S.J., Sandoval, J., Guan, G., and Coppens, I. (2011). Plasmodium salvages cholesterol internalized by LDL and synthesized de novo in the liver. Cellular microbiology 13, 569–586.

Lopes da Silva, M., Thieleke-Matos, C., Cabrita-Santos, L., Ramalho, J.S., Wavre-Shapton, S.T., Futter, C.E., Barral, D.C., and Seabra, M.C. (2012). The host endocytic pathway is essential for Plasmodium berghei late liver stage development. Traffic (Copenhagen, Denmark) 13, 1351–1363.

Mota, M.M., Pradel, G., Vanderberg, J.P., Hafalla, J.C., Frevert, U., Nussenzweig, R.S., Nussenzweig, V., and Rodriguez, A. (2001). Migration of Plasmodium sporozoites through cells before infection. Science 291, 141–144.

Niklaus, L., Agop-Nersesian, C., Schmuckli-Maurer, J., Wacker, R., Grunig, V., and Heussler, V.T. (2019). Deciphering host lysosome-mediated elimination of Plasmodium berghei liver stage parasites. Scientific reports 9, 7967.

Petersen, W., Stenzel, W., Silvie, O., Blanz, J., Saftig, P., Matuschewski, K., and Ingmundson, A. (2017). Sequestration of cholesterol within the host late endocytic pathway restricts liver-stage Plasmodium development. Molecular biology of the cell 28, 726–735.

Posfai, D., Maher, S.P., Roesch, C., Vantaux, A., Sylvester, K., Peneau, J., Popovici, J., Kyle, D.E., Witkowski, B., and Derbyshire, E.R. (2020). Plasmodium vivax Liver and Blood Stages Recruit the Druggable Host Membrane Channel Aquaporin-3. Cell Chem Biol.

Prado, M., Eickel, N., De Niz, M., Heitmann, A., Agop-Nersesian, C., Wacker, R., Schmuckli-Maurer, J., Caldelari, R., Janse, C.J., Khan, S.M., et al. (2015). Long-term live imaging reveals cytosolic immune responses of host hepatocytes against Plasmodium infection and parasite escape mechanisms. Autophagy 11, 1561–1579.

Prudencio, M., Rodrigues, C.D., Hannus, M., Martin, C., Real, E., Goncalves, L.A., Carret, C., Dorkin, R., Rohl, I., Jahn-Hoffmann, K., et al. (2008). Kinome-wide RNAi screen implicates at least 5 host hepatocyte kinases in Plasmodium sporozoite infection. PLoS Pathog 4, e1000201.

Raphemot, R., Toro-Moreno, M., Lu, K.Y., Posfai, D., and Derbyshire, E.R. (2019). Discovery of Druggable Host Factors Critical to Plasmodium Liver-Stage Infection. Cell Chem Biol 26, 1253–1262 e1255.

Real, E., Rodrigues, L., Cabal, G.G., Enguita, F.J., Mancio-Silva, L., Mello-Vieira, J., Beatty, W., Vera, I.M., Zuzarte-Luis, V., Figueira, T.N., et al. (2018). Plasmodium UIS3 sequesters host LC3 to avoid elimination by autophagy in hepatocytes. Nat Microbiol 3, 17–25.

Reed, N.A., Cai, D., Blasius, T.L., Jih, G.T., Meyhofer, E., Gaertig, J., and Verhey, K.J. (2006). Microtubule acetylation promotes kinesin-1 binding and transport. Curr Biol 16, 2166–2172.

Risco-Castillo, V., Topcu, S., Marinach, C., Manzoni, G., Bigorgne, A.E., Briquet, S., Baudin, X., Lebrun, M., Dubremetz, J.F., and Silvie, O. (2015). Malaria Sporozoites Traverse Host Cells within Transient Vacuoles. Cell host & microbe 18, 593–603.

Rodrigues, C.D., Hannus, M., Prudencio, M., Martin, C., Goncalves, L.A., Portugal, S., Epiphanio, S., Akinc, A., Hadwiger, P., Jahn-Hofmann, K., et al. (2008). Host scavenger receptor SR-BI plays a dual role in the establishment of malaria parasite liver infection. Cell host & microbe 4, 271–282.

Sanjana, N.E., Shalem, O., and Zhang, F. (2014). Improved vectors and genome-wide libraries for CRISPR screening. Nat Methods 11, 783–784.

Sciaky, N., Presley, J., Smith, C., Zaal, K.J., Cole, N., Moreira, J.E., Terasaki, M., Siggia, E., and Lippincott-Schwartz, J. (1997). Golgi tubule traffic and the effects of brefeldin A visualized in living cells. J Cell Biol 139, 1137–1155.

Shakibaei, M., and Frevert, U. (1996). Dual interaction of the malaria circumsporozoite protein with the low density lipoprotein receptor-related protein (LRP) and heparan sulfate proteoglycans. J Exp Med 184, 1699–1711.

Shalem, O., Sanjana, N.E., Hartenian, E., Shi, X., Scott, D.A., Mikkelson, T., Heckl, D., Ebert, B.L., Root, D.E., Doench, J.G., et al. (2014). Genome-scale CRISPR-Cas9 knockout screening in human cells. Science 343, 84–87.

Shortt, H.E., and Garnham, P.C. (1948). The pre-erythrocytic development of Plasmodium cynomolgi and Plasmodium vivax. Trans R Soc Trop Med Hyg 41, 785–795.

Silvie, O., Greco, C., Franetich, J.F., Dubart-Kupperschmitt, A., Hannoun, L., van Gemert, G.J., Sauerwein, R.W., Levy, S., Boucheix, C., Rubinstein, E., et al. (2006). Expression of human CD81 differently affects host cell susceptibility to malaria sporozoites depending on the Plasmodium species. Cellular microbiology 8, 1134–1146.

Sturm, A., Amino, R., van de Sand, C., Regen, T., Retzlaff, S., Rennenberg, A., Krueger, A., Pollok, J.M., Menard, R., and Heussler, V.T. (2006). Manipulation of host hepatocytes by the malaria parasite for delivery into liver sinusoids. Science (New York, NY) 313, 1287–1290.

Than, H., Pomicter, A.D., Yan, D., Beaver, L.P., Eiring, A.M., Heaton, W.L., Senina, A., Clair, P.M., Shacham, S., Mason, C.C., et al. (2020). Coordinated inhibition of nuclear export and Bcr-Abl1 selectively targets chronic myeloid leukemia stem cells. Leukemia.

Tran, T.M., Guha, R., Portugal, S., Skinner, J., Ongoiba, A., Bhardwaj, J., Jones, M., Moebius, J., Venepally, P., Doumbo, S., et al. (2019). A Molecular Signature in Blood Reveals a Role for p53 in Regulating Malaria-Induced Inflammation. Immunity 51, 750–765 e710.

Vanderberg, J.P. (1981). Plasmodium berghei exoerythrocytic forms develop only in the liver. Trans R Soc Trop Med Hyg 75, 904–905.

Vijayan, K., Cestari, I., Mast, F.D., Glennon, E.K.K., McDermott, S.M., Kain, H.S., Brokaw, A.M., Aitchison, J.D., Stuart, K., and Kaushansky, A. (2019). Plasmodium Secretion Induces Hepatocyte Lysosome Exocytosis and Promotes Parasite Entry. iScience 21, 603–611.

Wacker, R., Eickel, N., Schmuckli-Maurer, J., Annoura, T., Niklaus, L., Khan, S.M., Guan, J.L., and Heussler, V.T. (2017). LC3-association with the parasitophorous vacuole membrane of Plasmodium berghei liver stages follows a noncanonical autophagy pathway. Cellular microbiology 19.

Warncke, J.D., and Beck, H.P. (2019). Host Cytoskeleton Remodeling throughout the Blood Stages of Plasmodium falciparum. Microbiol Mol Biol Rev 83.

White, M.T., Walker, P., Karl, S., Hetzel, M.W., Freeman, T., Waltmann, A., Laman, M., Robinson, L.J., Ghani, A., and Mueller, I. (2018). Mathematical modelling of the impact of expanding levels of malaria control interventions on Plasmodium vivax. Nature communications 9, 3300.

Wiese, C., and Zheng, Y. (2000). A new function for the gamma-tubulin ring complex as a microtubule minus-end cap. Nat Cell Biol 2, 358–364.

Zhu, X., and Kaverina, I. (2011). Quantification of asymmetric microtubule nucleation at subcellular structures. Methods Mol Biol 777, 235–244.

Zhu, X., and Kaverina, I. (2013). Golgi as an MTOC: making microtubules for its own good. Histochem Cell Biol 140, 361–367.

## References

Geldenhuys, W.J., Leeper, T.C., and Carroll, R.T. (2014). mitoNEET as a novel drug target for mitochondrial dysfunction. Drug Discov Today 19, 1601–1606.

Kim, H.C., Jo, Y.J., Kim, N.H., and Namgoong, S. (2015). Small molecule inhibitor of formin homology 2 domains (SMIFH2) reveals the roles of the formin family of proteins in spindle assembly and asymmetric division in mouse oocytes. PLoS One 10, e0123438.

Luedtke, D.A., Su, Y., Liu, S., Edwards, H., Wang, Y., Lin, H., Taub, J.W., and Ge, Y. (2018). Inhibition of XPO1 enhances cell death induced by ABT-199 in acute myeloid leukaemia via Mcl-1. J Cell Mol Med 22, 6099–6111.

Martinez-Navarro, F.J., Martinez-Morcillo, F.J., Lopez-Munoz, A., Pardo-Sanchez, I., Martinez-Menchon, T., Corbalan-Velez, R., Cayuela, M.L., Perez-Oliva, A.B., Garcia-Moreno, D., and Mulero, V. (2020). The vitamin B6-regulated enzymes PYGL and G6PD fuel NADPH oxidases to promote skin inflammation. Dev Comp Immunol 108, 103666.

Miyazawa, M., Bogdan, A.R., and Tsuji, Y. (2019). Perturbation of Iron Metabolism by Cisplatin through Inhibition of Iron Regulatory Protein 2. Cell Chem Biol 26, 85–97 e84.

Pulvino, M., Liang, Y., Oleksyn, D., DeRan, M., Van Pelt, E., Shapiro, J., Sanz, I., Chen, L., and Zhao, J. (2012). Inhibition of proliferation and survival of diffuse large B-cell lymphoma cells by a small-molecule inhibitor of the ubiquitin-conjugating enzyme Ubc13-Uev1A. Blood 120, 1668–1677.

Santos, C., Fleury, L., Rodriguez, F., Markus, J., Berkes, D., Daich, A., Ausseil, F., Baudoin-Dehoux, C., Ballereau, S., and Genisson, Y. (2015). The CERT antagonist HPA-12: first practical synthesis and individual binding evaluation of the four stereoisomers. Bioorg Med Chem 23, 2004–2009.

Ye, H., Zhao, Q., Huang, Y., Wang, L., Liu, H., Wang, C., Dai, D., Xu, L., Ye, M., and Duan, S. (2014). Metaanalysis of low density lipoprotein receptor (LDLR) rs2228671 polymorphism and coronary heart disease. Biomed Res Int 2014, 564940.

